# FET fusion proteins reshape splicing factor networks to drive oncogenic alternative splicing

**DOI:** 10.64898/2026.08.28.746988

**Authors:** Loïc Ongena, Eva Lucarelli, Laurence Dubois, Jonathan Bruyr, Lianghao Mao, Florian H. Geyer, Florencia Cidre-Aranaz, Thomas G. P. Grünewald, Ashok Kumar Jayavelu, Yiyun Zhang, Akshay Bhinge, Didier Vertommen, Franck Dequiedt

## Abstract

Gene fusions involving the FET gene family (*FUS*, *EWSR1*, and *TAF15*) act as drivers of numerous cancer entities. The resulting chimeric proteins are widely viewed as aberrant transcriptional regulators that promote malignant transformation through chromatin and enhancer reprogramming. Here, we show that FET fusion oncoproteins also function as regulators of alternative splicing across multiple sarcoma entities. Transcriptomic analyses revealed extensive but largely non-overlapping splicing programs driven by the EWSR1::FLI1, EWSR1::WT1, EWSR1::ATF1 and FUS::DDIT3 fusions that converged on common oncogenic functions. Fusion-dependent splicing regulation was mechanistically separable from canonical transcriptional activity and was associated with extensive remodeling of cooperative RNA-binding protein (RBP) assemblies on target transcripts. Despite regulating distinct exons, different FET fusions engaged highly similar RBP interaction networks, consistent with a conserved mode of splicing regulation. Transcriptome-wide mapping of RBP occupancy revealed extensive reorganization of local RNA regulatory landscapes following fusion depletion. The requirement of RNA for FET fusion condensate formation, together with the inability of condensation-defective mutants to restore splicing regulation, further supported a role for higher-order assemblies in fusion-dependent alternative splicing (AS) control. Fusion-driven splicing programs stratified Ewing sarcoma patients independently of established clinical covariates, thereby underscoring their clinical relevance. AS of *TFDP1* emerged as a common fusion-regulated splicing event required for sarcoma cell fitness and therapeutically actionable using antisense oligonucleotides. Together, our findings establish AS regulation as a conserved function of FET fusion oncoproteins that is mechanistically separable from their canonical transcriptional activity. More broadly, they support a model in which oncogenic fusion proteins can drive malignant phenotypes through large-scale remodeling of RNA regulatory networks.

## Main

Gene fusions generated by chromosomal rearrangements represent major oncogenic drivers across multiple cancer entities, particularly sarcomas and hematological malignancies [1,2]. A substantial fraction of these rearrangements involves transcription factors (TFs), generating oncogenic chimeric transcription factors (OCTFs) that profoundly rewire transcriptional and epigenetic programs as well as cellular identity [3,4]. As master regulators of gene expression, OCTFs therefore provide powerful models to investigate how aberrant regulation of gene expression drives malignant transformation.

The FET gene family, comprising *FUS*, *EWSR1* and *TAF15*, encodes multifunctional RNA-binding proteins (RBPs) involved in multiple aspects of RNA metabolism [5]. FET genes are recurrently rearranged across diverse sarcoma and leukemia entities, where they frequently act as pathognomonic oncogenic drivers. These rearrangements generate OCTFs that almost systematically fuse the N-terminal intrinsically disordered low-complexity domain (LCD) of FET proteins to the DNA-binding domain (DBD) of various unrelated TFs. Owing to this conserved domain organization, FET fusion oncoproteins have been extensively characterized as aberrant transcriptional regulators that profoundly reshape epigenomic landscapes [6,7]. Mechanistically, FET fusions recruit chromatin remodeling and transcriptional regulatory complexes to ectopic genomic regions, thereby promoting *de novo* enhancer activation, disrupting higher-order chromatin architecture, and ultimately altering transcriptional outputs [8–12].

The prion-like LCD retained in FET fusions mediates multivalent molecular interactions and promotes the assembly of dynamic biomolecular condensates through liquid-liquid phase separation (LLPS) [13–15]. These properties critically depend on aromatic residues, particularly tyrosines distributed throughout the LCD, and are thought to facilitate the local concentration of fusion-associated cofactors at specific genomic loci, thereby potentiating regulatory activity [16–19]. Importantly, interactomic studies have shown that FET fusions engage in extensive protein interaction networks enriched in RBPs and RNA-processing factors, suggesting functions extending beyond transcriptional control [20,21].

Alternative splicing (AS) constitutes a major layer of post-transcriptional gene regulation that substantially expands transcriptomic and proteomic diversity [22,23]. Its dysregulation contributes to multiple hallmarks of cancer, and recurrent alterations affecting splicing regulators are observed across diverse cancer entities [24]. Splicing decisions are governed by combinatorial assemblies of RBPs on *cis*-regulatory elements within pre-mRNAs, which collectively control spliceosome recruitment and exon definition [22,25]. Increasing evidence further indicates that these ribonucleoprotein (RNP) assemblies can organize into condensate-like structures that spatially coordinate splicing outcomes [26]. Given their propensity to form condensates and interact with numerous splicing-associated RBPs, FET fusion oncoproteins are therefore ideally positioned to influence AS regulation. However, despite several reports implicating FET fusions in splicing regulation, the extent, underlying mechanisms, and biological significance of FET fusion-dependent AS regulation remain poorly understood [20,27–30]. Moreover, most studies have so far focused on EWSR1::FLI1, the archetypal fusion defining Ewing sarcoma, leaving the broader impact of FET fusion oncoproteins on transcriptome-wide isoform regulation largely unexplored.

Here, we systematically investigated the role of FET fusion oncoproteins in AS across four prototypical sarcoma models driven by EWSR1::FLI1 (Ewing sarcoma, EwS), EWSR1::WT1 (desmoplastic small round cell tumor, DSRCT), EWSR1::ATF1 (clear cell sarcoma, CCS), and FUS::DDIT3 (myxoid liposarcoma, MLPS). Using integrated transcriptomic, proteomic, biochemical, and functional approaches, we demonstrate that FET fusions establish conserved oncogenic splicing programs that are largely independent of their canonical transcriptional activities. Mechanistically, our findings support a model in which FET fusions associate with pre-mRNAs through interactions with RBPs and reshape local splicing regulatory networks in an RNA- and condensate-dependent manner. Finally, we identify clinically relevant fusion-dependent splicing events associated with patient outcome, thereby establishing AS regulation as a previously underappreciated function of FET fusions in the promotion of malignant processes.

## Results

### FET fusions establish conserved alternative splicing programs across sarcoma subtypes

To define the AS programs controlled by FET fusion oncoproteins, we analyzed RNA sequencing (RNA-seq) datasets from models of EwS, DSRCT, CCS, and MLPS, in the presence and absence of the corresponding fusion (EWSR1::FLI1, EWSR1::WT1, EWSR1::ATF1 and FUS::DDIT3, respectively). We reanalyzed published datasets for EwS and DSRCT models [31,32] and generated complementary RNA-seq profiles for CCS and MLPS cells. The integrated dataset comprised 12 sarcoma cell lines analyzed under matched control and fusion-depleted conditions, including four EwS (shA673-1C, TC-71, MHH-ES-1 and SK-N-MC), two DSRCT (BER and JN-DSRCT-1), and three CCS (MP-CCS-SY, KAS and SU-CCS-1) and MLPS (DL-221, MLS-402 and MLS-1765) cell lines (**Figure 1A**). Principal component analysis (PCA) demonstrated robust clustering according to both sarcoma entity and knockdown conditions (**Figure S1A,B**).

**Figure 1.**
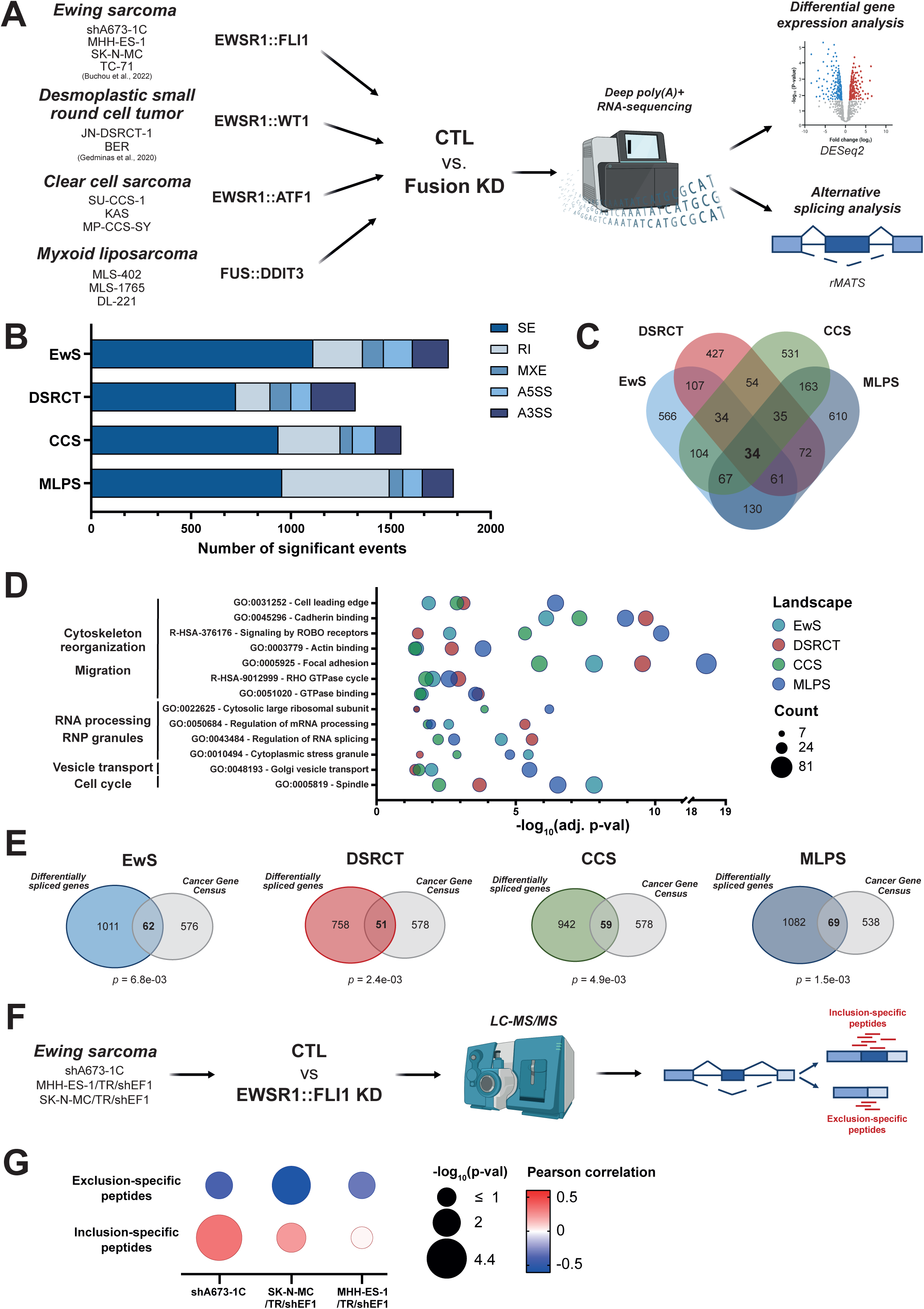
FET fusions establish conserved alternative splicing programs across sarcoma subtypes. (**A**) Schematic workflow of FET fusion-dependent transcriptome analysis. (**B**) Alternative splicing landscapes under FET fusion control across the Ewing sarcoma (EwS), desmoplastic small round cell tumor (DSRCT), clear cell sarcoma (CCS) and myxoid liposarcoma (MLPS) models. SE: skipped exon; RI: retained intron; MXE: mutually exclusive exon; A5SS: alternative 5’ splice site; A3SS: alternative 3’ splice site. (**C**) Overlap of differentially spliced genes across the four splicing landscapes. (**D**) Shared gene ontology and pathway enrichment across the four sarcoma models. P-values were adjusted using the Benjamini-Hochberg method. (**E**) Overlap of differentially spliced genes with the COSMIC Cancer Gene Census set. *p*: overlap p-value (Fisher’s exact test). (**F**) Schematic workflow for the analysis of the EWSR1::FLI1-dependent proteome by tandem mass spectrometry. (**G**) Correlations between the fold-changes in the abundance of peptides mapping to exclusion- or inclusion-specific junctions and the ΔPSI levels in three EwS cell lines.

Consistent with their established functions as oncogenic transcriptional regulators, FET fusions controlled the expression of several thousand genes across sarcoma models (**Figure S1C**). Differential gene expression analyses recapitulated established fusion-specific transcriptional programs (**Figure S1D**). In EwS and DSRCT models, FET fusions promoted cell cycle-associated transcriptional programs while repressing pathways linked to cell migration. In CCS cells, EWSR1::ATF1 suppressed subsets of MITF-dependent melanocytic differentiation pathways, whereas FUS::DDIT3 in MLPS altered transcriptional programs associated with lipid metabolism and neuronal signaling. Despite distinct fusion-specific transcriptional signatures, recurrent enrichment of pathways related to cell cycle regulation and cell migration was observed across sarcoma models, suggesting convergence toward shared oncogenic programs.

Given the emerging evidence linking FET fusions to RNA processing, we next investigated whether these oncoproteins establish AS programs in their respective sarcoma cells. We quantified differential exon inclusion across the five major classes of AS events, including skipped exons (SE), retained introns (RI), mutually exclusive exons (MXE), and alternative 5’ and 3’ splice site usage (A5SS and A3SS). Transcriptome-wide splicing analysis using rMATS revealed extensive AS regulation in every model, with thousands of significant splicing events detected following fusion depletion (**Figure S1E**). Representative AS events were independently validated by RT-PCR, confirming the reproducibility of the RNA-seq analysis (**Figure S1F**).

Pairwise comparison of the percent spliced in (PSI) values for shared SE events revealed strong and highly significant correlations between related cell lines of the same tumor type (**Figure S1G**). We therefore defined high-confidence sarcoma-specific “splicing landscapes” composed of recurrent AS events displaying consistent inclusion patterns across related cell lines. Each landscape comprised more than one thousand reproducible AS events, with SEs representing the predominant class across all sarcoma models, followed by RIs, alternative splice sites and MXEs (**Figure 1B, Supplementary Table S1**). Most differentially spliced genes (DSGs) contained a single regulated event, supporting exon-specific regulation rather than generalized splicing disruption (**Figure S1H**). Because splice site selection is strongly influenced by local *cis*-acting features, we next investigated various properties of fusion-regulated AS events. Regulated exons displayed remarkably similar characteristics across all sarcoma landscapes, including weaker splice sites (**Figure S1I**), elevated GC content surrounding exon-intron junctions (**Figure S1J**), enrichment in RNA G-quadruplex (**Figure S1K**), and preferential downstream positioning within transcripts relative to background events (**Figure S1L**). In addition, retained introns and exons involved in A5SS exhibited increased sequence length (**Figure S1M,N**). Together, these findings indicate that FET fusion-dependent AS programs preferentially target exons with recurrent non-random *cis*-regulatory properties, suggesting the existence of shared determinants underlying splice site selection across FET fusion sarcomas.

Comparison of the four splicing landscapes revealed limited overlap at the level of individual AS events. Only nine events were common to all landscapes, the majority of which were skipped exons. These recurrent events affected transcripts involved in translation initiation (*EIF4A2*), protein ubiquitination (*WSB1* and *ADRM1*), phospholipid metabolism (*SCARB1* and *PI4KB*), protein transport (*AP2M1*), and cell cycle regulation (*TFDP1*). Similarly, only 34 DSGs were common to all landscapes, indicating that FET fusions primarily regulate distinct repertoires of splicing events and transcripts (**Figure 1C**). Functional enrichment analysis revealed biological processes associated with fusion-regulated DSGs within each landscape (**Figure S1O**). Surprisingly, despite the limited overlap between individual AS events or DSGs, several functional categories were recurrently enriched across all four landscapes, including pathways linked to RNA metabolism, vesicle transport, cytoskeletal remodeling, migration, and cell cycle regulation (**Figure 1D**). Therefore, while FET fusions regulate relatively few common genes or splicing events, they control common biological functions through alternative splicing control. Cancer Gene Census annotations were significantly enriched in DSGs from each landscape, indicating that distinct FET fusions broadly reshape the splicing programs of cancer-associated genes (**Figure 1E**).

Because alternative splicing can alter transcript stability through nonsense-mediated decay (NMD), we next assessed the predicted coding potential of fusion-regulated isoforms. Most regulated events were classified as protein-coding rather than NMD-sensitive transcripts, based on two complementary approaches using transcript annotations and overlap with experimentally determined AS events associated with NMD (**Figure S1P, Q**) [33]. This suggests that FET fusion-dependent AS programs predominantly generate stable transcript isoforms. We next investigated whether these transcript variants produced detectable protein isoforms by performing proteomic analyses in three EwS cell lines following EWSR1::FLI1 depletion. Peptides corresponding to alternatively spliced isoforms were classified as inclusion- or exclusion-specific (**Figure 1F**). Changes in inclusion- and exclusion-specific peptide abundance generally correlated with ΔPSI values measured by RNA-seq across the three cell lines (**Figure 1G**). These results indicate that a substantial fraction of FET fusion-dependent splicing changes is reflected at the protein level, supporting their potential to alter the proteomic output of fusion-driven sarcomas.

### Splicing control by FET fusions is independent of transcriptional regulation

We next investigated a potential link between fusion-dependent splicing and transcriptional regulation. DSGs displayed non-significant overlap with differentially expressed genes across all sarcoma models, indicating that splicing and transcriptional regulation by FET fusions primarily affect distinct gene populations (**Figure 2A**). Consistent with this observation, ChIP-Seq analysis revealed limited and non-significant overlaps between genes associated with fusion binding and DSGs in each sarcoma subtype (**Figure S2A-D**). Moreover, more than 70% of fusion-regulated AS events lack detectable fusion occupancy within a 100 kb window surrounding the regulated region (**Figure 2B**). Together, these findings indicate that FET fusion-dependent AS regulation operates largely independently of genomic binding and transcriptional regulation.

**Figure 2.**
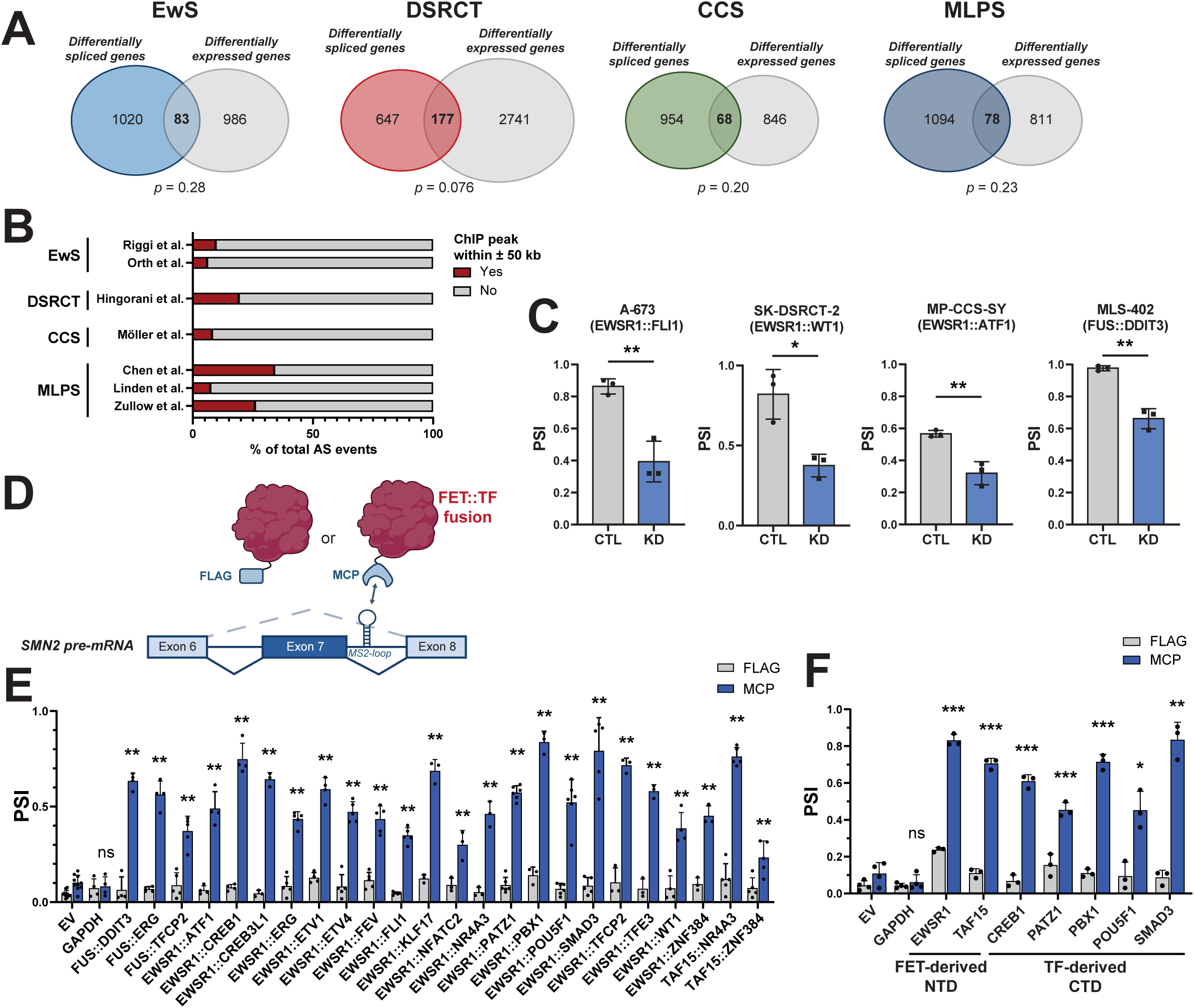
Splicing control by FET fusions is independent of transcriptional regulation. (**A**) Overlap between differentially expressed and differentially spliced genes. *p*: overlap p-value (Fisher’s exact test). (**B**) Fraction of splicing events that overlap a ChIP-seq peak within ±50 kb. (**C**) Differential splicing of the *TFDP1* minigene following fusion knockdown across the four sarcoma models. *: p-value < 0.05; **: p-value < 0.01 (two-sided unpaired t-test). (**D**) Schematic representation of the MS2-tethering splicing assay in which the *SMN2* reporter minigene is co-transfected with FET fusions tagged with either FLAG or the MS2-coat protein (MCP), which enables its specific recruitment to MS2-binding sites (MS2-bs) on the reporter pre-mRNA. (**E**) Exon inclusion of the *SMN2* reporter minigene upon transfection of empty vector (EV) and GAPDH controls, and a panel of 24 different FET fusions. (**F**) Exon inclusion of the *SMN2* reporter minigene upon transfection of empty vector (EV) and GAPDH controls, and fusion N-terminal and C-terminal domains. PSI: percentage spliced in. Results depicted represent at least 3 independent biological replicates. *: p-value < 0.05; **: p-value < 0.01 (FDR-adjusted Wilcoxon–Mann–Whitney test).

To confirm this observation and determine whether fusion-dependent splicing regulation required endogenous genomic and chromatin contexts, we generated splicing reporter minigenes for three fusion-regulated SE events affecting *EIF4A2*, *TFDP1* and *MACROH2A1*. All reporters recapitulated endogenous splicing patterns and were responsive to fusion depletion across multiple sarcoma models (**Figure S2E, S1F**). Notably, the *TFDP1* minigene, corresponding to a conserved event shared across all splicing landscapes, consistently displayed robust fusion-dependent regulation (**Figure 2C, S2E**). These findings indicate that splicing regulation by FET fusions is independent of promoter identity, chromatin states and higher-order genome organization.

Based on these findings, we next asked whether FET fusions could directly regulate splicing through physical association with pre-mRNA. To also determine whether this activity extends beyond the four sarcoma models analyzed above, we used an MS2-based tethering assay to recruit a panel of 24 structurally diverse FET fusions to an alternatively spliced *SMN2* reporter minigene (**Figure 2D**). Strikingly, all 24 fusions promoted exon inclusion when tethered to the reporter RNA, whereas untethered constructs failed to significantly alter splicing patterns (**Figure 2E**). Comparable results were obtained using an independent *MAPT* reporter minigene, although with lower overall amplitudes (**Figure S2G, H**). Because these fusions contain highly related FET-derived N-terminal domains fused to otherwise unrelated TF-derived C-terminal regions, we initially expected their splicing activity to primarily depend on the FET moiety. Surprisingly, however, both the FET-derived N-terminal domain and TF-derived C-terminal domain were individually sufficient to promote exon inclusion upon RNA recruitment (**Figure 2G**). Together, these findings demonstrate that FET fusion oncoproteins possess intrinsic splicing regulatory activity that depends on physical association with pre-mRNA and involves contributions from both the FET-derived and TF-derived regions.

### FET fusions regulate splicing through RBP interactions

Because recruitment to pre-mRNA was required for fusion-dependent splicing regulation, we next investigated whether FET fusion could associate with RNA. Using Orthogonal Organic Phase Separation (OOPS) assays [34] in representative models of EwS, DSRCT, CCS, and MLPS, we observed that all four fusions were specifically recovered within RNP complexes under combined ultraviolet (UV) light and disuccinimidyl glutarate (DiSG) crosslinking conditions, which preserve both direct RNA-protein and protein-protein interactions (**Figure 3A, B**). By contrast, no direct RNA interaction was detected following UV-only crosslinking, suggesting that FET fusions associate with RNA indirectly, most likely via protein-protein interactions with RBPs.

**Figure 3.**
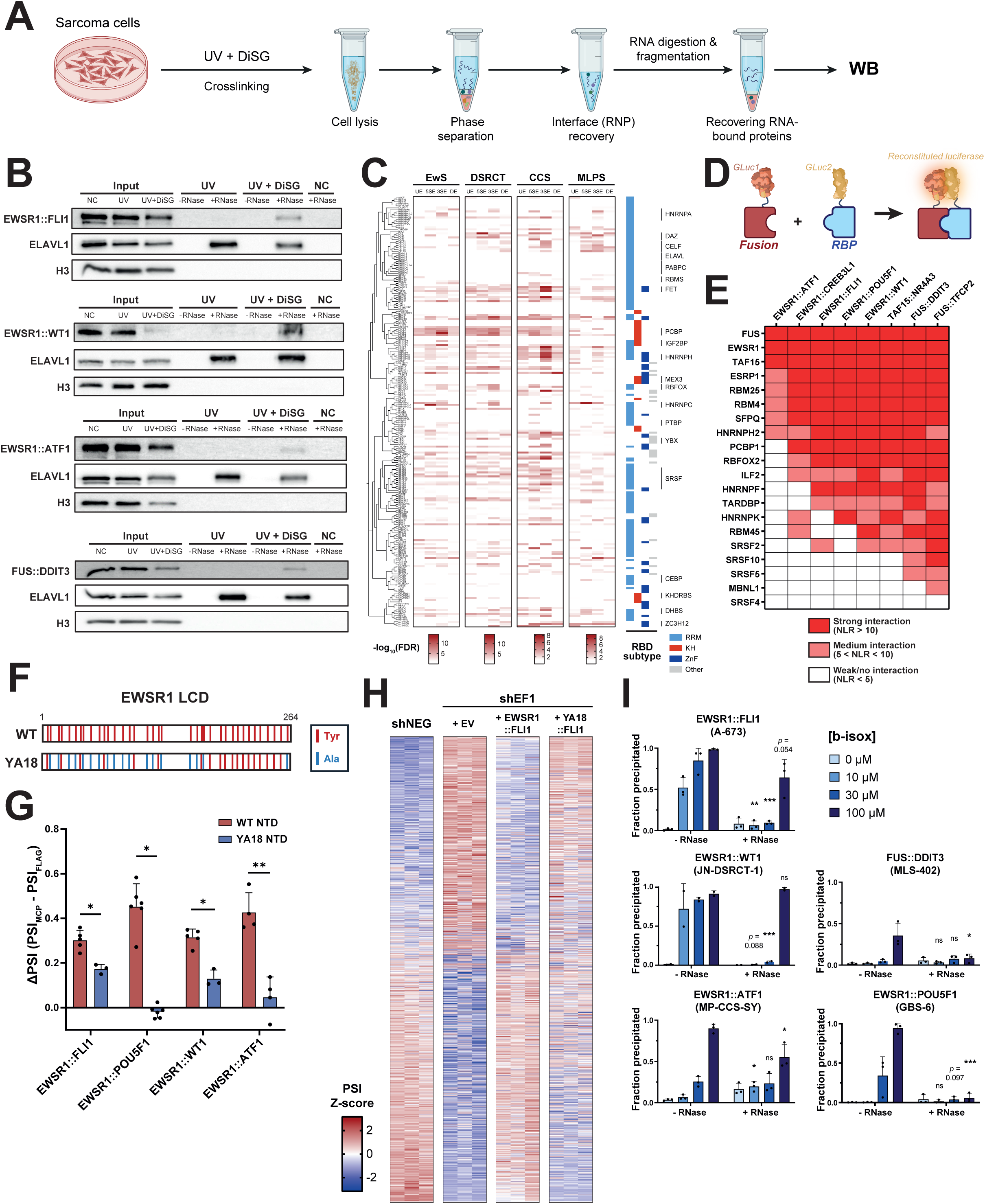
FET fusions regulate splicing through RBP interactions. (**A**) Experimental workflow of the OOPS assay. DiSG: disuccinimidyl glutarate; RNP: ribonucleoprotein; WB: western blot. (**B**) Representative western blots of protein fractions resulting from the OOPS assay in EwS, DSRCT, CCS and MLPS cells. +/- RNase designates experimental conditions in which RNA was (+) or was not (−) degraded following interface recovery. NC: non-crosslinked. (**C**) Motif enrichment analysis around fusion-regulated splicing events. Significance results of the analysis were aggregated for each RBP in the four splicing junctions for each landscape. RBPs were clustered based on sequence homology (CLUSTALW alignment) and grouped by subfamily. RNA-binding domain (RBD) subtypes are depicted on the right side of the heatmap. P-values inferior to 0.05 (FDR-adjusted one-sided Wilcoxon– Mann–Whitney test) are not colored. (**D**) Principle of the *Gaussia* luciferase protein complementation assay (GPCA) in which luciferase activity is reconstituted and luminescence is emitted upon interaction of protein partners tagged with two complementary moieties of the *Gaussia princeps* luciferase (GLuc1 and GLuc2). (**E**) Mean normalized luminescence ratios (NLR) of the interactions between twenty RBPs and eight representative FET fusions spanning diverse families. Results depicted represent at least 3 independent biological replicates. (**F**) Representation of the EWSR1 low-complexity domain (LCD) with tyrosine mutations. (**G**) Splicing activity of fusion tyrosine mutants represented as the net differential exon inclusion of the *SMN2* reporter minigene upon transfection of FLAG/MCP-tagged constructs of wild-type (WT) or the YA18 fusion mutants. *: p-value < 0.05; **: p-value < 0.01 (Wilcoxon–Mann– Whitney test). (**H**) Exon inclusion levels upon fusion knockdown and re-expression of WT and mutant fusions in Ewing sarcoma cells. Heatmap represents inclusion levels across four conditions ranked from lowest to highest relative inclusion in the control condition: control (shNEG), knockdown (shEF1+EV), knockdown rescued by EWSR1::FLI1, and knockdown rescued by the YA18::FLI1 mutant. Depicted are z-score transformed exon inclusion levels of events rescued by the wild-type fusion (FDR < 0.05). Each column represents an independent biological replicate. (**I**) RNA-dependent precipitation of endogenous FET fusions with increasing concentrations of biotinylated isoxazole (b-isox). ns: not significant; *: p-value < 0.05; **: p-value < 0.01; ***: p-value < 0.001 (two-sided unpaired t-test).

To identify RBPs potentially involved in fusion-dependent AS regulation, we performed systematic motif enrichment analyses within each splicing landscape using a sliding-window approach centered on exon-intron junctions (**Figure S3A**), generating positional maps for known RBP motifs associated with each SE dataset. Robust enrichment patterns for numerous RBPs from diverse RNA-binding domain families were observed in every landscape (**Figure 3C**). Strikingly, many RBP motifs were consistently enriched across all four landscapes despite limited overlap between regulated SEs themselves, suggesting the recruitment of partially shared RBP repertoires by distinct FET fusions. However, only limited positional differences were observed around regulated SEs, and no consistent pattern emerged when comparing included and excluded events (**Figure S3B**). The predicted binding of many RBPs was further supported by orthogonal computational approaches and enrichment of experimentally validated CLIP-seq binding sites (**Figure S3C**). Altogether, we identified a compendium of 51 RBPs from diverse families (highlighted in **Figure S3C**) that fulfilled multiple criteria, including: (i) recurrent enrichment across motif and/or CLIP-based analyses in all landscapes, (ii) established roles in pre-mRNA splicing, (iii) known associations with oncogenic processes, and (iv) detectable expression in most sarcoma models. Consistent with the sequence properties of fusion-regulated SE identified above, many of these RBPs preferentially recognize G- or GC-rich sequences and have been implicated in rG4 recognition [35].

A simple explanation for fusion-dependent AS regulation would be altered expression of splicing factors following fusion depletion. However, no enrichment of RNA processing or splicing-related pathways was detected in any differential expression dataset (**Figure S1D**). Moreover, few of the 51 candidate RBPs displayed transcriptional or proteomic dysregulation following fusion knockdown, and these changes were not consistent across sarcoma models (**Figure S3D, E**). Together, these findings indicate that fusion-dependent AS regulation is unlikely to arise from indirect changes in splicing factor abundance. We therefore hypothesized that FET fusions regulate AS through physical interactions with RBPs. Supporting this possibility, more than half (28/51) of the enriched RBPs had previously been identified in published interactomes of EWSR1::FLI1 or FUS::DDIT3, including the splicing-associated partners FUS, HNRNPA1, HNRNPC, HNRNPH1, HNRNPK, NONO, SFPQ and SRSF9 interacting with both fusions [20,21]. To directly test FET fusions/RBPs interactions, we quantitatively assessed binary protein-protein interactions between representative FET fusions and a panel of 20 RBPs using *Gaussia* luciferase protein complementation assays (GPCA) (**Figure 3D**). Several RBPs, including EWSR1, FUS, TAF15, ESRP1, HNRNPH2, RBM4, RBM25 and SFPQ, displayed robust interactions with all tested fusions (**Figure 3E**). By contrast, no reproducible interaction was detected with SRSF4, SRSF5, SRSF10 or MBNL1, whereas other RBPs interacted with most, but not all FET fusions. These findings provide a mechanistic explanation for how FET fusion oncoproteins are recruited to RNA despite lacking intrinsic RNA-binding activity.

Because condensate-like assemblies have been implicated in the spatial organization of RNA processing and splicing [26], we next investigated whether the condensate-forming properties of FET fusions are required for their splicing activity. To address this question, we substituted 18 tyrosine residues with alanines in the EWSR1 LCD of EWSR1::FLI1, EWSR1::POU5F1, EWSR1::WT1 and EWSR1::ATF1, generating the YA18::FLI1, YA18::POU5F1, YA18::WT1 and YA18::ATF1 mutants, respectively (**Figure 3F**). Using biotinylated isoxazole (b-isox) precipitation assays as a proxy for condensation propensity, we observed that the YA18 mutations severely impaired condensation properties of all four FET fusions while largely preserving their nuclear localization (**Figure S3F, G**). When tested in a tethering assay, all YA18-mutated fusions exhibited markedly reduced splicing activity compared with their wild-type counterpart (**Figure 3G**). Consistent with these observations, re-analysis of previously published RNA-seq data [36] revealed that, unlike wild-type EWSR1::FLI1, the condensation-defective YA18 mutant failed to restore most fusion-dependent AS events following EWSR1::FLI1 depletion (**Figure 3H**). These findings indicate that condensation-associated properties of the FET LCD are important for fusion-dependent splicing regulation.

Because RNA can contribute to the formation of RNP condensates, we next investigated whether RNA is involved in the biomolecular assembly properties of FET fusions. As expected, we observed dose-dependent b-isox precipitation of all four endogenous FET fusions in their respective sarcoma cells (**Figure 3I**). Remarkably, RNase A treatment markedly altered precipitation profiles across all models, indicating that RNA contributes to the assembly or stability of fusion-associated condensate-like structures. Together, these findings support a model in which FET fusions regulate splicing through multivalent interactions with RBPs within RNA-dependent condensate-associated networks.

### FET fusions disrupt splicing factor networks

Because splicing regulation relies on cooperative interactions between RBPs assembled on pre-mRNAs, we next investigated whether FET fusions alter the organization of local splicing networks. We first examined whether the 51 core fusion-associated RBPs identified above tend to co-occur around regulated exons using local motif co-occurrence analyses centered on exon-intron junctions. Pairwise comparisons of motif frequencies revealed complex and highly interconnected landscape-specific co-occurrence networks (**Figure 4A, B**). In all cases, the vast majority of RBPs were connected to at least one partner, supporting widespread motif co-occurrence around fusion-regulated exons. Nodes corresponding to members of the HNRNPH, RBM4, and FET protein families consistently displayed high betweenness centrality across all four networks, suggesting central roles in the assembly of local RBP hubs associate with FET fusions. Interestingly, many interactions displayed strong positional specificity and were preferentially enriched either upstream or downstream of cassette exons. These findings suggest that fusion-regulated splicing decisions are coordinated through the targeted assembly of distinct local RBP subnetworks rather than through the isolated activity of individual factors. To further explore the cooperative potential of these RBPs, we reconstructed protein-protein interaction networks involving the 51 fusion-associated splicing factors using curated interactomes and the HiNT database [37]. Strikingly, nearly all core RBPs (49/51) interacted with at least one additional fusion-associated factor, forming a densely interconnected interactome (**Figure 4C**). Together, these observations indicate that FET fusion-associated RBPs possess extensive cooperative potential at both the RNA-binding and protein-interaction levels.

**Figure 4.**
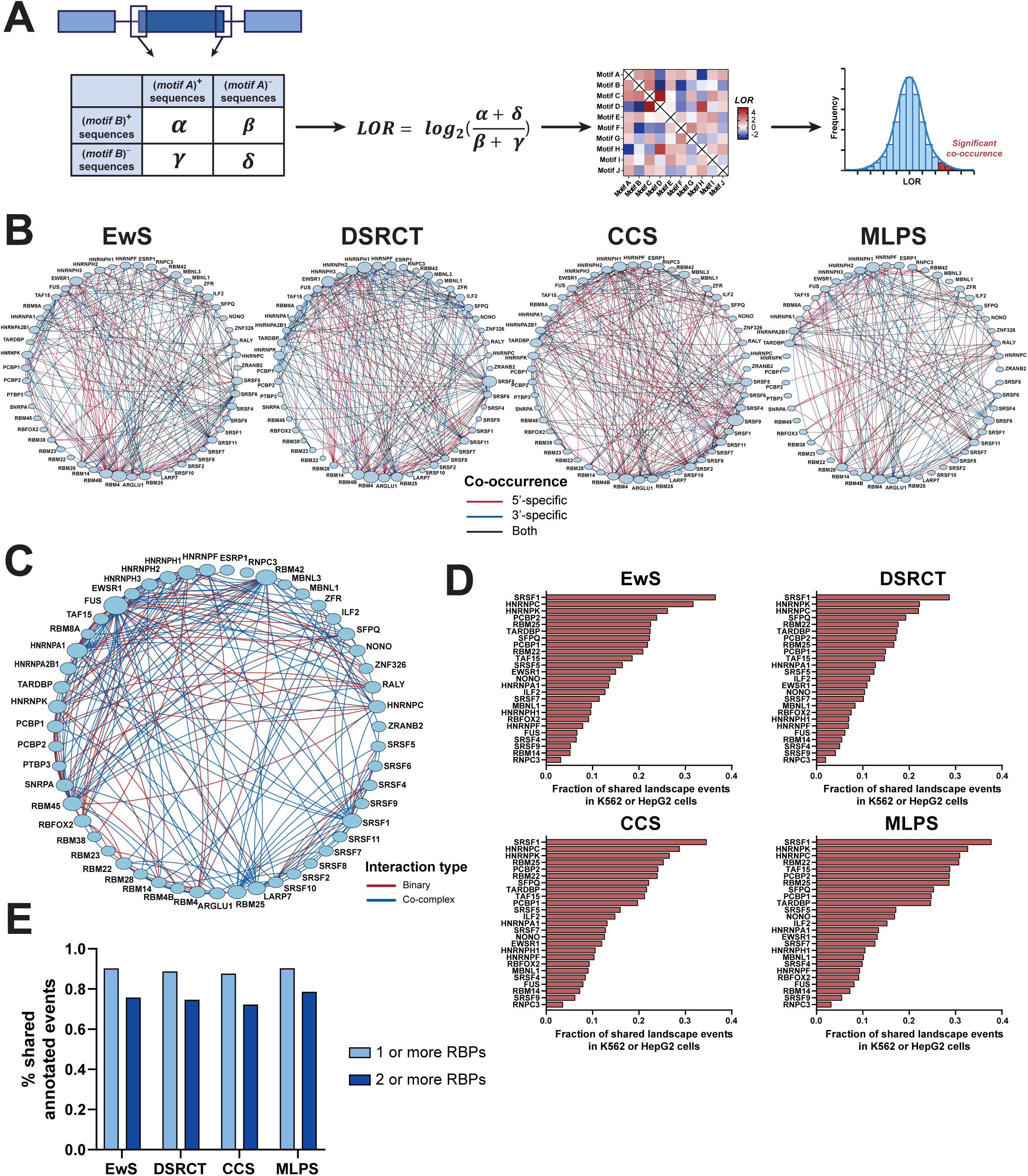
FET fusions disrupt splicing factor networks. (**A**) Workflow of the motif co-occurrence computational analysis. Pairwise comparisons of motif frequencies within 5’ and 3’ exon-intron junctions of differentially skipped exons are used to compute the log-transformed odds-ratio (LOR) score which serves as a proxy for motif co-occurrence. (**B**) Motif co-occurrence between partner RBPs across the four splicing landscapes. LOR scores were aggregated per RBP and displayed as a network in which edges represent significant motif co-occurrence between two RBPs (nodes). These co-occurrence patterns were either specific to the 5’ (red) or 3’ (blue) junction or common in both junctions (black). Node size is proportional to the number of edges connected to it. (**C**) Protein-protein interaction network of partner splicing factors, with binary and co-complex interactions represented as red and blue edges, respectively. Node size is proportional to the number of edges connected to it. (**D**) Fraction of co-regulated splicing events dependent on both fusion and RBP expression, regardless of directionality, based on RNA-seq data in K562 and/or HepG2 from the ENCODE project. (**E**) Fraction of annotated events shared by at least one (light blue) or at least two (dark blue) different partner RBPs.

We next asked whether candidate RBPs functionally co-regulate fusion-dependent AS events. Analysis of ENCODE RNA-seq datasets [38] following depletion of 26 core RBPs in K562 and HepG2 cells revealed substantial overlaps between RBP- and fusion-regulated splicing profiles (**Figure 4D**). In particular, events regulated by SRSF1, HNRNPC, HNRNPK, RBM22, RBM25, PCBP2 and SFPQ overlapped with 15-35% of events within individual fusion-dependent splicing landscapes. Overall, nearly 90% of fusion-regulated events were also controlled by at least one core RBP, whereas more than 70% were co-regulated by at least two distinct RBPs across all landscapes (**Figure 4E**). Because AS regulation is highly cell type-specific, we hypothesized that ENCODE datasets derived from unrelated cellular contexts underestimate the extent of co-regulation in sarcoma cells. To address this possibility, we performed RNA-seq analyses following depletion of RBFOX2 in EwS, DSRCT, CCS, and MLPS models (**Figure S4A**). In sarcoma cells, the overlap between RBFOX2- and fusion-regulated events increased approximately threefold relative to ENCODE cell lines (**Figure S4B**). Moreover, RBFOX2 depletion produced non-random regulatory relationships with fusion-dependent splicing programs, with globally opposite effects in EwS and DSRCT cells but concordant effects in CCS and MLPS cells. Together, these findings support a model in which fusion-dependent AS is governed by a coordinated network of multiple RBPs, rather than by individual splicing factors.

Based on these observations, we hypothesized that FET fusions directly perturb RBP occupancy on target transcripts. To investigate this possibility, we performed exonuclease-assisted mapping of protein-RNA interactions (ePRINT), a transcriptome-wide footprinting approach that identifies RBP-protected RNA regions (**Figure 5A**) [39]. We applied ePRINT following doxycycline-inducible depletion of EWSR1::FLI1 in the EwS shA673-1C cell line. In total, 47,831 and 52,613 reproducible peaks were identified in control and fusion-depleted conditions, respectively. Most peaks exhibited sharp 5′ boundaries consistent with exonuclease arrest at protein-protected regions and a biphasic pattern, suggestive of adjacent RBP binding events (**Figure S5A**). Peaks were predominantly localized within introns, with additional enrichment in coding regions and 3′UTRs (**Figure S5B**). Moreover, more than 90% of peaks overlapped CLIP-seq binding sites for at least one RBP, supporting the specificity of the dataset (**Figure S5C**). Together, these observations establish ePRINT as a robust approach to profile RBP occupancy landscapes in EwS cells. Differential binding analyses revealed extensive remodeling of RBP footprints following EWSR1::FLI1 knockdown. Although 19,861 peaks were shared between fusion-expressing and knocked down cells, 31,418 peaks were preferentially detected following fusion knockdown, whereas 26,933 peaks were enriched in fusion-expressing cells (**Figure S5D**). Restricting analyses to fusion-regulated AS regions revealed that approximately half of regulated events overlapped at least one ePRINT peak, and more than 80% of these events displayed peak changes upon fusion depletion, with 6,909 and 8,081 peaks specific to the control and KD conditions, respectively (**Figure 5B**). No consistent relationship emerged between peak remodeling patterns, positional distribution around SE and changes in exon inclusion, suggesting that splicing outcomes are unlikely to be determined by simple positional effects of RBP binding (**Figure S5E**). Remarkably, motifs corresponding to 45 of the 51 fusion-associated core RBPs were significantly enriched within differential peaks (**Figure 5C**). Binding sites derived from corresponding ENCODE eCLIP datasets were similarly enriched within shifting peaks for most core RBPs (**Figure S5F**). Together, these findings support a model in which FET fusions reshape local RBP occupancy landscapes on target pre-mRNAs, thereby altering the regulatory networks that govern splice site selection.

**Figure 5.**
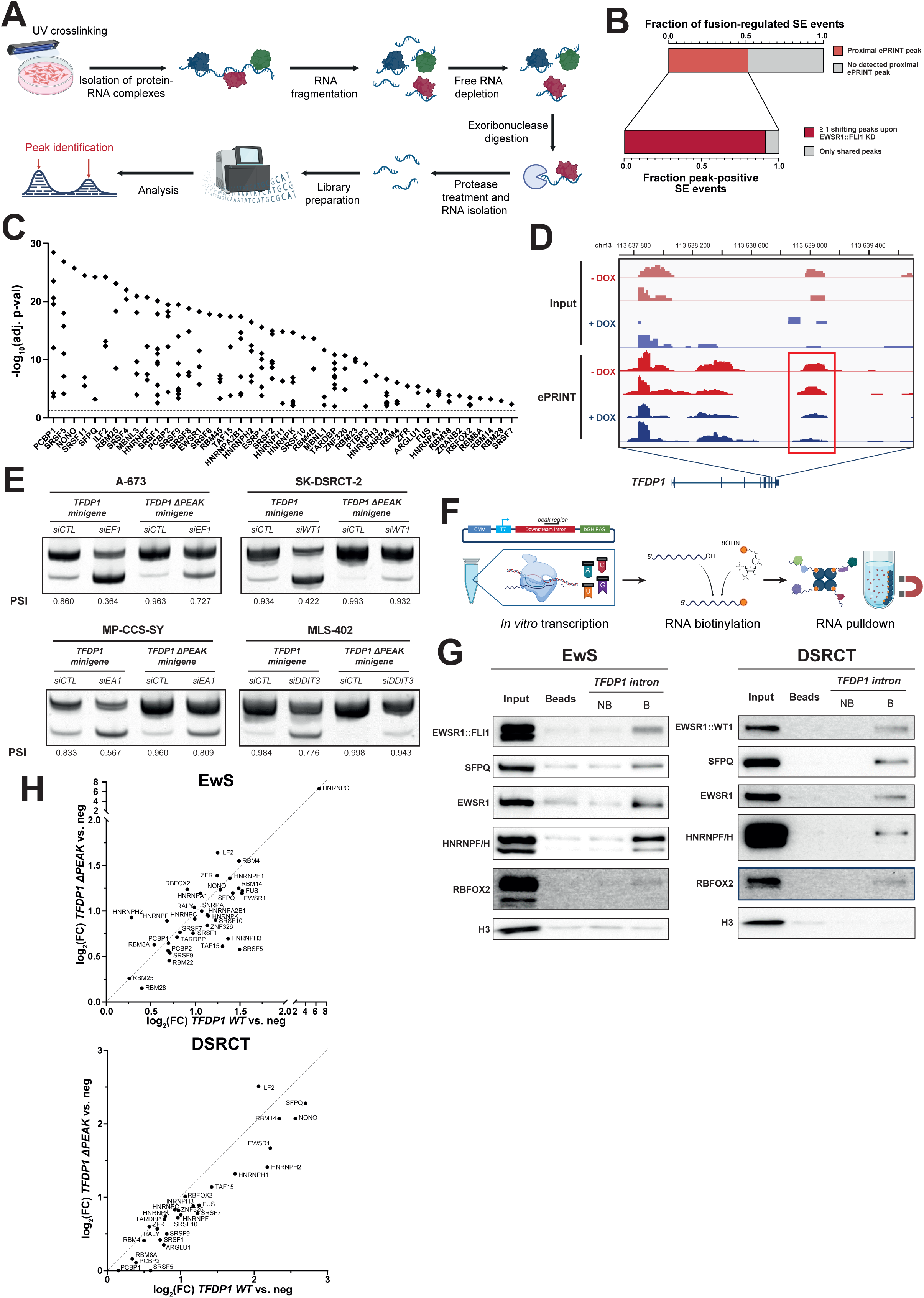
RBPs are broadly redistributed around fusion-regulated splicing events. (**A**) Workflow of the ePRINT experiment. (**B**) Fraction of EWSR1::FLI1-dependent skipped exon (SE) events overlapping ePRINT peaks in shA673-1C cells and the fraction of these events with one or more shifting peaks upon fusion knockdown. (**C**) Results of the AME (MEME suite) motif enrichment analysis of shifting vs. shared ePRINT peaks around fusion-regulated events. Enriched motifs were aggregated per RBP. (**D**) ePRINT track profiles downstream of the *TFDP1* cassette exon. (**E**) Representative RT-PCR results of cassette exon inclusion levels of the WT and ΔPEAK *TFDP1* minigene transcripts upon fusion knockdown in four sarcoma models. (**F**) Workflow of the RNA pulldown experiment. (**G**) Representative western blots of the recovered proteins bound to the biotinylated *TFDP1* intron RNA with EwS and DSRCT cell lysates. B: pulldown with biotinylated RNA; NB: pulldown with non-biotinylated RNA (control). Beads: pulldown with streptavidin beads only. H3 serves as a negative control. (**H**) Correlation between the log-transformed fold changes of the relative abundance of proteins bound to the wild-type and mutated *TFDP1* downstream intron based on proteomic analysis of RNA pulldowns with EwS and DSRCT cell lysates.

Among EWSR1::FLI1-regulated SE, *TFDP1* displayed prominent ePRINT footprints surrounding the skipped exon (exon 11), including a strong downstream intronic peak that was markedly reduced following EWSR1::FLI1 depletion (**Figure 5D**). *TFDP1* exon 11 being one of the conserved AS events shared across all fusion-dependent splicing landscapes, we used it as a model to investigate fusion-mediated RBP redistribution. Deletion of the ePRINT peak region from the *TFDP1* minigene strongly reduced fusion-dependent splicing regulation across all sarcoma models, demonstrating that this intronic region contains *cis*-regulatory elements required for the control of *TFDP1* AS by FET fusions (**Figure 5E**). To identify RBPs binding to this region, we performed RNA pulldown assays using an RNA fragment centered on the *TFDP1* intronic ePRINT peak (**Figure 5F**). Validating this approach, the intronic RNA recovered EWSR1::FLI1 and EWSR1::WT1 fusions from EwS and DSRCT lysates, respectively, together with several of the 51 core RBPs implicated in fusion-dependent AS regulation, including EWSR1, SFPQ, RBFOX2 and HNRNPF/H proteins (**Figure 5G**). Tandem mass spectrometry analyses further revealed strong enrichment of candidate RBPs on the *TFDP1* intronic RNA (**Supplementary Table S2**). Comparative analysis between wild-type and ΔPEAK RNAs, showed reduced binding of most core RBPs upon deletion of the peak region in both EwS and DSRCT (**Figure 5H**). Together, these findings indicate that FET fusions regulate AS by reshaping cooperative RBP assemblies on target transcripts through specific *cis*-regulatory RNA elements.

### Fusion-dependent splicing programs are associated with patient outcome and identify *TFDP1* alternative splicing as a functional vulnerability

To investigate the clinical relevance of fusion-dependent AS programs in EwS, we analyzed RNA-seq and clinical data from 57 EwS patients from the International Cancer Genome Consortium (ICGC) cohort (**Figure S6A**). PSI values were assigned for all quantifiable AS events using a unified computational pipeline. Only events confidently detected in more than 80% of patients were retained for downstream analyses. Unsupervised hierarchical clustering based on exon inclusion profiles separated patients into two major groups of comparable size (**Figure S6B**). Strikingly, these clusters displayed significantly different overall survival, with cluster A patients exhibiting markedly poorer outcomes (**Figure 6A**). Importantly, multivariate Cox regression analyses confirmed that patient cluster assignment remained a prognostic factor independent from sex, metastatic status and age at diagnosis (**Table S1**). Together, these findings indicate that global splicing signatures are strongly associated with patient survival in EwS.

**Figure 6.**
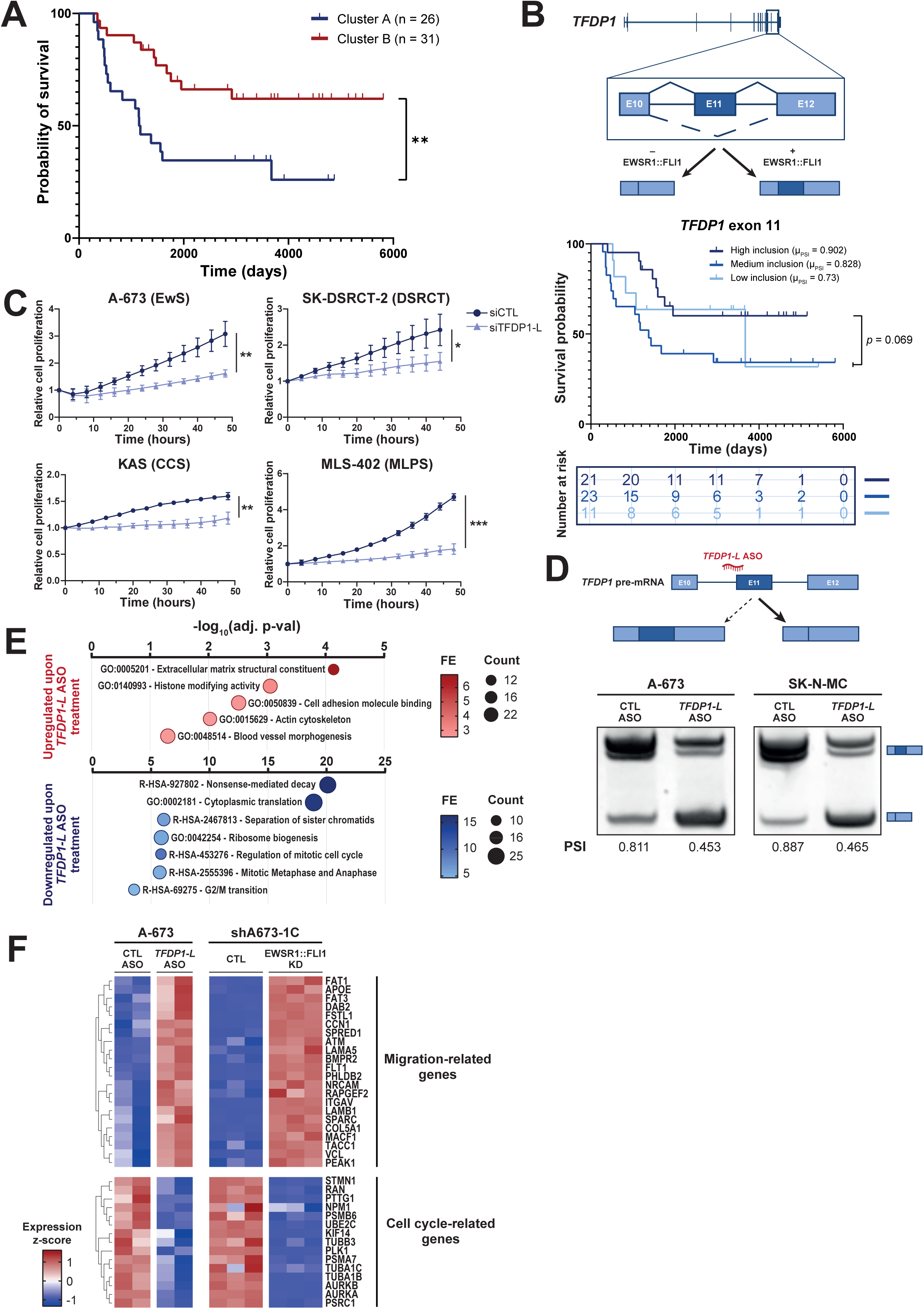
Fusion-dependent splicing programs are associated with patient outcome and identify TFDP1 alternative splicing as a functional vulnerability. (**A**) Kaplan-Meier estimation curve of the two main patient clusters defined by hierarchical clustering of Ewing sarcoma patient splicing profiles. n: number of patients. **: p-value < 0.01 (log-rank Mantel-Cox test). (**B**) Schematic representation and Kaplan-Meier estimation curve of the survival-associated EWSR1::FLI1-dependent AS events in the TFDP1 gene. p: p-value (log-rank Mantel-Cox test). (**C**) Proliferation curves of A-673, SK-DSRCT-2, KAS, and MLS-402 in control (siCTL) and knockdown of the long TFDP1 isoform (siTFDP1-L) conditions. Cell count was normalized to T0 and measured over 44 h. *: p-value < 0.05; **: p-value < 0.01; ***: p-value < 0.001 (two-way ANOVA test between conditions) (**D**) Top. Schematic representation of antisense oligonucleotide (ASO)-mediated redirection of TFDP1 splicing. Bottom. Representative RT-PCR result of isoform switch 72 h after ASO transfection in the A-673 (200 nM) and SK-N-MC (100 nM) EwS cells. (**E**) Gene ontology and pathway enrichment analysis of differentially regulated genes following ASO transfection in A-673 cells (200 nM, 72 h). FE: fold-enrichment. (**F**) Representative gene sets related to migration and cell cycle cellular processes in A-673 following ASO treatment (200 nM, 72 h) and in shA673-1C following fusion knockdown. Expression counts were z-score normalized per row for each dataset.

We next specifically examined the prognostic relevance of EWSR1::FLI1-regulated AS events. EwS landscape SEs with sufficient junction coverage in at least half of patients were retained for analysis, and PSI values were stratified into low-, intermediate- and high-inclusion groups using k-means clustering. Among 710 evaluable SE events, 138 (19.4%) were significantly associated with patient survival, indicating that a substantial fraction of fusion-regulated AS events has prognostic value. Both positively and negatively regulated EWSR1::FLI1-dependent exons were associated with prognosis, with no systematic relationship between exon inclusion direction and patient outcome (**Figure S6C**). Consistent with previous reports, increased expression of the short metastasis-associated *ADD3* isoform correlated with poorer survival [30]. Notably, increased inclusion of *TFDP1* exon 11, a conserved fusion-promoted event detected in EwS, DSRCT, CCS and MLPS landscapes, was associated with improved survival (**Figure 6B**). These findings highlight *TFDP1* exon 11 as a clinically relevant, conserved fusion-dependent splicing event for further functional investigation.

Because FET fusions consistently promote inclusion of *TFDP1* exon 11 and expression of a long *TFDP1-L* isoform, we next investigated the role of the corresponding protein in sarcoma cells. Selective depletion of TFDP1-L using isoform-specific siRNAs consistently impaired proliferation and cell viability across multiple sarcoma models (**Figure 6C**, **S6D, E**), indicating that the fusion-associated isoform contributes to sarcoma cell fitness. We next asked whether selective redirection of *TFDP1* splicing could reproduce the phenotypic consequences of fusion depletion and provide a therapeutically actionable means of targeting this fusion-dependent splicing event. To specifically assess the functional contribution of *TFDP1* splicing, we designed a splice-switching antisense oligonucleotide (ASO) to promote exon 11 skipping. This approach reproduced the *TFDP1* isoform switch observed following fusion knockdown, while preserving the other regulatory activities of the fusion (**Figure 6D, S6F**). To assess the functional consequences of this switch, we performed RNA-seq analysis of EwS cells following treatment with the *TFDP1-L*-targeting ASO and identified 382 differentially expressed genes (adjusted p-value < 0.05). Gene ontology and pathway enrichment analyses showed that genes upregulated following ASO treatment were associated with cytoskeletal and extracellular matrix reorganization, whereas downregulated genes were enriched for processes related to translation, cell cycle progression and mitosis (**Figure 6E**). This transcriptional response closely mirrors the signature associated with fusion knockdown, suggesting that a component of the oncogenic program driven by EWSR1::FLI1 is mediated through its regulation of *TFDP1* splicing. Consistent with this interpretation, 194 out of the 382 genes differentially expressed following *TFDP1* isoform switching were similarly dysregulated following EWSR1::FLI1 knockdown (adjusted p-value < 0.05, concordant direction), including genes associated with cell cycle progression and migration (**Figure 6F**). Together, these findings identify *TFDP1* as a conserved functional target of FET fusion-dependent AS and suggest that fusion-driven regulation of *TFDP1* splicing contributes to the transcriptional and phenotypic programs that sustain sarcoma cells. The ability to selectively redirect this splicing event further supports pathogenic AS programs as potentially actionable downstream outputs of FET fusion activity.

## Discussion

FET fusion oncoproteins have so far largely been viewed as aberrant transcription factors that drive sarcomagenesis through chromatin and enhancer reprogramming. Our findings support a broader model in which FET fusions also function as RNA regulatory organizers that reshape cooperative splicing factor networks. Despite controlling largely non-overlapping splicing events, distinct FET fusions converge on common oncogenic programs linked to proliferation, migration, cytoskeletal organization, and RNA metabolism. Importantly, multiple observations indicate that this splicing activity is mechanistically separable from their canonical transcriptional regulation. These include the limited overlap between differentially spliced and transcriptionally regulated genes, the absence of fusion occupancy at most regulated exons, and the preservation of fusion-dependent splicing outside endogenous chromatin contexts. Alternative splicing therefore emerges not as a secondary consequence of fusion-driven transcriptional rewiring, but as a crucial layer of FET fusion biology.

FET fusions do not behave as classical splicing factors. Rather than recognizing specific RNA motifs, they appear to regulate splicing through reorganization of cooperative RBP assemblies on target transcripts via multivalent protein-protein interactions. This is notably consistent with prior reports that both RBFOX2 and EWSR1 physically and functionally cooperate with EWSR1::FLI1 in splicing regulation within different contexts [30,40]. Our model shifts the regulatory unit from the individual splice site to the broader RBP network assembled on pre-mRNAs. As distinct FET fusions target largely different exons, perturbations of dynamic and cell-specific RBP complexes could generate heterogeneous exon-level outputs while still producing convergent oncogenic states, consistent with models of coordinated post-transcriptional RNA regulons.

Our findings further suggest that this RNA splicing activity represents a neomorphic property created by coupling the FET LCD to heterologous transcription factor-derived regions. Despite the structural diversity of their C-terminal partners, all tested FET fusions retained remarkably similar splicing activity upon RNA recruitment, arguing for a unified underlying mechanism. At the same time, both the FET-derived N-terminal region and the TF-derived C-terminal domains independently modulated splicing when tethered to pre-mRNA, suggesting that fusion-dependent splicing regulation arises from cooperation between a conserved FET interaction platform and fusion-specific regulatory properties contributed by the TF partner. This architecture parallels emerging models in which intrinsically disordered domains drive higher-order regulatory organization across multiple layers of gene expression [41,42]. Taken together, these observations raise the possibility that RNA-centered regulatory activities may represent a previously underappreciated property of oncogenic fusion transcription factors with low-complexity activation domains.

The RNA-dependent nature of fusion assemblies, together with the loss of splicing activity in condensation-defective mutants, suggest that FET fusions regulate splicing within dynamic RNA-associated assemblies rather than through isolated molecular interactions. These findings indicate that the condensation-associated properties of FET fusions contribute not only to transcriptional and chromatin regulation, but also to the organization of RBP assemblies controlling splice site selection. In this context, condensate formation may not simply concentrate regulatory factors but instead stabilize cooperative RBP networks and reshape their local organization on target transcripts. These results extend current models of FET fusion condensates beyond chromatin-associated functions and suggest that analogous assembly principles operate at the level of RNA processing.

Our study identifies *TFDP1* splicing as a conserved functional target across the four sarcoma models. Consistent with prior studies [43–45], the *TFDP1-L* isoform, favored in the presence of FET fusions, was associated with increased cell proliferation and contributed to sarcoma cell fitness. Conversely, promoting *TFDP1* exon 11 skipping induced transcriptional changes associated with cell migration and angiogenesis and partially recapitulated the transcriptional response to fusion depletion. This finding is consistent with the phenotypic plasticity of EwS cells, in which highly proliferative EWSR1::FLI1-high states often remain less invasive than more dedifferentiated fusion-low states associated with metastasis and therapeutic resistance [46]. *TFDP1* splicing may therefore contribute to the stabilization of specific fusion-associated cellular states and serve as a marker of this plasticity. Importantly, the ability to redirect *TFDP1* splicing using ASOs in a sarcoma model further demonstrates that this fusion-dependent splicing event can be therapeutically manipulated downstream of FET fusion activity.

More broadly, our findings identify fusion-dependent splicing programs as biologically important and therapeutically actionable features of FET fusion sarcomas. Global splicing signatures stratified EwS patients independently of established clinical covariates, while a substantial fraction of fusion-regulated events carried prognostic value. These observations suggest that fusion-dependent AS programs capture biologically meaningful dimensions of tumor state that are not fully reflected by conventional transcriptional or clinical classifications. Given the large number of OCTFs retaining intrinsically disordered activation domains [47], it is tempting to speculate that RNA-centered regulatory activities analogous to those described here may extend far beyond FET fusion sarcomas. Our findings therefore support a broader view of fusion oncoprotein biology in which RNA processing represents an additional layer of oncogenic regulation alongside chromatin and transcriptional control. Remodeling of cooperative RNA regulatory networks may thus represent a broader principle by which oncogenic fusion proteins reshape cellular state.

## Acknowledgments

We thank all members of the laboratory of Gene Expression and Cancer for insightful discussions and inputs. We thank the GIGA Genomics and Bioinformatics platforms for their technical support and helpful assistance. We acknowledge use of the GIGA high performance computing cluster for conducting the study reported in this paper. This work was supported by the University of Liège (ULiège), the Belgian National Fund for Scientific Research (F.R.S-FNRS), Télévie, the GIGA-Doctoral School for Health Sciences, the Fondation contre le Cancer, the Roi Baudouin Foundation and the Fondation Léon Fredericq. The laboratory of F.C.A. and T.G.P.G. acknowledges funding by the Dr. Leopold and Carmen Ellinger Foundation, the Henrik-Kreibohm Foundation, and the European Union (ERC, CANCER-HARAKIRI, 101122595). Views and opinions expressed are however those of the authors only and do not necessarily reflect those of the European Union or the European Research Council (ERC). F.H.G. was supported by scholarships of the German Cancer Aid and the German Academic Scholarship Foundation.

## Author contributions

L.O. and F.D. conceptualization; L.O., E.L. and F.D. methodology; L.O., E.L., J.B., L.D., L.M., Y.Z. experimental investigation; L.O. bioinformatic analyses; A.K.J., A.B., D.V., F.H.G., F.C.A., T.G.P.G., and F.D. resources; L.O. and F.D. writing; F.D. supervision and funding acquisition.

## Methods

### Cell lines and cell culture

HeLa, HEK-293T, SU-CCS-1 and A-673 cells were purchased from the American Type Culture Collection (ATCC), and DL-221 cells from the MDACC Cytogenetics and Cell Authentication Core (MD Anderson Cancer Center, USA). The EwS cell line shA673-1C was a gift from Dr. Olivier Delattre (Institut Curie, France). The EwS cell lines SK-N-MC/TR/shEF1, MHH-ES-1/TR/shEF1, and TC-71/TR/shEF1 were described previously [48]. Parental EwS cell lines were obtained from the following repositories: SK-N-MC and MHH-ES-1 from DSMZ (German Collection of Microorganisms and Cell Cultures), and TC-71 from the Children’s Oncology Group (COG) childhood cancer repository. The DSRCT cell lines JN-DSRCT-1 and SK-DSRCT-2 were kindly provided by Dr. Mikiko Aoki (Fukuoka University, Fukuoka, Japan) and by Dr. Marc Ladanyi (Memorial Sloan Kettering Cancer Center, New York, NY, USA), respectively. MLS-402 and MLS-1765 cells were gifts from Dr. David Braig (University of Freiburg, Germany) and Dr. Pierre Åman (Sahlgrenska Cancer Center, Sweden), respectively. MP-CCS-SY cells were gifts from Dr. Hiroshi Moritake (University of Miyazaki, Japan), and KAS and GBS-6 cells from Dr. Takuro Nakamura (Japanese Foundation for Cancer Research, Japan).

All cells were cultured in a humidified incubator at 37°C with 5% CO_2_. HeLa, HEK-293T, A-673, DL-221 and SK-DSRCT-2 cells were maintained in Dulbecco’s Modified Eagle’s Medium (DMEM, Biowest) with 10% Fetal Bovine Serum (FBS, Greiner). MLS-402, MLS-1765, KAS, MP-CCS-SY, TC-71, SK-N-MC, MHH-ES-1 and SU-CCS-1 cells were maintained in RPMI 1640 medium (Biowest) with 10% FBS, and JN-DSRCT-1 cells in Advanced DMEM/F-12 (ThermoFisher) with 10% FBS and L-glutamine (ThermoFisher). shA673-1C cells were additionally cultured with 10 µg/mL blasticidin and 200 µg/mL zeocin (InvivoGen), and SK-N-MC/TR/shEF1, MHH-ES-1/TR/shEF1, and TC-71/TR/shEF1 cells with 1 µg/mL puromycin (InvivoGen), to maintain the corresponding shRNA constructs. shRNA-mediated knockdown (KD) was induced with 1 µg/mL doxycycline (dox) for 5-7 days, refreshed every 48 h. All experiments used cells between passages 2 and 15. Cell lines were routinely tested for mycoplasma contamination with the MycoAlert™ Mycoplasma Detection Kit (Lonza) by the GIGA-Viral Vectors platform (University of Liège).

### Plasmids and cloning

Open reading frames (ORFs) encoding proteins of interest were amplified by high-fidelity PCR (Phusion™ DNA polymerase, New England Biolabs) using primers flanked by Gateway attB sequences, then inserted into pDONR223 vectors by BP cloning (Invitrogen, Gateway system). All ORFs contained an ATG start codon and lacked a STOP codon to allow subcloning into multiple tag configurations. Fusion ORFs were generated by overlap-extension PCR using complementary flanking primers spanning the fusion junction. Validated pDONR223 constructs were transferred by LR cloning (Invitrogen) into pDEST1899-FLAG (N-terminal 3xFLAG), pDEST-MCP (N-terminal MS2 coat protein-3xFLAG) or pDEST-GLucN1/N2 (N-terminal Gaussia princeps luciferase fragments) expression vectors. The EWSR1-derived tyrosine mutants YS7 and YS37 were generated by gene synthesis (Genscript). The YA18 mutant was a gift from Dr. Kevin Lee (Hong Kong University of Science and Technology, China). Coding sequences of *NFATC2*, *TFE3*, *NR4A3*, *KLF17* and *TARDBP* were amplified from Addgene plasmid templates (**Supplementary Table S3**). All other ORFs were retrieved or derived from the human ORFeome v7.1/v8.1 collection (Center for Cancer Systems Biology, Dana-Farber Cancer Institute).

All vectors were transformed into *Escherichia coli* DH5α and cultured in lysogeny broth with the appropriate antibiotic. Plasmids were extracted with the NucleoSpin Plasmid kit (Macherey-Nagel), eluted in nuclease-free water, and quantified with a NanoDrop spectrophotometer (NanoPhotometer® N60, Implen). All constructs were verified by Sanger or whole-plasmid Nanopore sequencing (Eurofins Genomics). Cloning primers, manufactured by Eurogentec (Liège, Belgium), are listed in **Supplementary Table S3**.

### Chemicals

Biotinylated isoxazole (b-isox, Focus Biomolecules) was dissolved in DMSO as 10 mM aliquots and stored at −20°C for up to one month. Doxycycline hyclate (dox, Sigma-Aldrich) was dissolved in DMSO to 100 µg/mL and stored at −20°C. Disuccinimidyl glutarate (DiSG, Proteochem) was dissolved in DMSO to 0.25 M and diluted to the working concentration in cold PBS immediately before use.

### Transfection

Plasmid DNA was transfected into HeLa and HEK-293T cells with polyethyleneimine (PEI) 40K (2:1 PEI/DNA mass ratio), and into sarcoma cells with jetPRIME® (Polyplus), per the manufacturers’ protocols. Transient knockdowns were achieved using small interfering RNAs (siRNAs, Eurogentec, Liège, Belgium; sequences in **Supplementary Table S3**) transfected with Lipofectamine RNAiMAX (Invitrogen). Expression and knockdown efficiency were assessed by western blotting 48-72 h after transfection. For previously published datasets, RNA was collected 48 h after siRNA transfection in DSRCT cells, 96 h after transfection in TC-71, SK-N-MC and MHH-ES-1 cells, and after 7 days of dox treatment in shA673-1C cells. 2’-O-methoxyethyl-phosphorothioated antisense oligonucleotides (Integrated DNA Technologies; sequences in **Supplementary Table S3**) targeting the 3’ splice site upstream of *TFDP1* exon 11 were transfected into cells with Lipofectamine RNAiMAX (Invitrogen). Splice switch efficiency was assessed by RT-PCR and western blotting 72 h after transfection.

### RNA extraction, reverse transcription and polymerase chain reaction

Total RNA was extracted from PBS-washed cells with the NucleoSpin RNA kit (Macherey-Nagel), eluted in RNase-free water and quantified by NanoDrop. RNA (500 ng-1 µg) was reverse-transcribed with random hexamers using the FastGene Scriptase II cDNA kit (Nippon Genetics). PCR was performed on cDNA with Taq polymerase (ThermoFisher), with annealing temperatures (50-60°C) and elongation times adjusted to primer melting temperature and amplicon length. Primers for endogenous and minigene splicing assays are listed in **Supplementary Table S3**. PCR products were resolved on 12% polyacrylamide gels, stained for 30 min with GelStar fluorescent dye (1:10,000, Lonza) in Tris-Borate-EDTA buffer (10.5 g/L Tris, 5.7 g/L boric acid, 2 mM EDTA, pH 8), washed twice with dH2O, and imaged under UV light on an ImageQuant LAS 4000 (GE Healthcare Life Sciences). Band intensities were quantified with ImageQuantTL software, and percent spliced-in (PSI) values were calculated as the ratio of the longer isoform band intensity to the sum of both isoforms.

For RNA-seq samples, total RNA quality was assessed with a BioAnalyzer (Agilent), with RNA integrity numbers exceeding 9 for all samples. Libraries were prepared with the TruSeq Stranded mRNA kit (Illumina, poly(A) selection) and sequenced paired-end (150 bp, >50M reads/sample) on a NovaSeq™ 6000 v1.5 platform (Illumina) by the GIGA-Genomics platform (University of Liège).

### Western blotting

Cell lysates were boiled for 5 min in Laemmli buffer (62.5 mM Tris-HCl, 10% glycerol, 2% SDS, 3% β-mercaptoethanol, 0.0003% bromophenol blue) and resolved by SDS-PAGE in Tris-Glycine-SDS buffer. Proteins were transferred onto 0.2 µm nitrocellulose membranes (ThermoFisher Scientific) in Tris-Glycine buffer with 10% methanol, blocked for 1 h in TBS-T with 5% non-fat milk, and incubated overnight at 4°C with primary antibodies in TBS-T with 5% milk or 4% BSA. Membranes were then incubated with HRP-linked secondary antibodies for 1 h, washed three times, and developed by chemiluminescence on an ImageQuant LAS 4000 system (GE Healthcare Life Sciences) using a homemade substrate solution (Solution A: 50 mg Luminol sodium salt [Merck] in 200 mL 0.1 M Tris-HCl, pH 8.6; Solution B: 11 mg p-Coumaric acid [Merck] in 10 mL DMSO; Solution C: 35% H2O2; mixed 1:0.1:0.0003). Band intensities were quantified with ImageQuantTL software. Primary and secondary antibodies are listed in **Supplementary Table S3**.

### Minigene assays

Minigenes of the FET fusion splicing targets *EIF4A2*, *MACROH2A1* and *TFDP1*, along with *TFDP1* mutant constructs, were amplified by high-fidelity PCR from sarcoma cell genomic DNA and cloned into a HindIII-linearized pcDNA3.1 vector by NEBuilder HiFi DNA assembly (New England Biolabs). Because the genomic region spanning the upstream-to-downstream exon of MACROH2A1 was too long to amplify as one fragment, exon-containing segments with 500 bp intronic extensions were ligated sequentially. A *TFDP1* deletion mutant, lacking an internal sequence element of the downstream intron, was generated by overlap-extension PCR.

Sarcoma cells (A-673, SK-DSRCT-2, MP-CCS-SY and MLS-402) were plated in 6-well plates, transfected at ∼70% confluence with control or fusion-targeting siRNAs, and transfected 24 h later with 500 ng of the corresponding minigene plasmid. Cells were collected 48 h after plasmid transfection, washed with PBS, and split for western blotting (to confirm KD efficacy) and RNA extraction followed by RT-PCR (to assess exon inclusion). PCR used a forward primer targeting the T7 promoter of pcDNA3.1 and a reverse primer hybridizing to the downstream exon of the target event. PCR products were resolved on 12% acrylamide gels as described above.

### MS2-tethering reporter assays

The previously reported *SMN2-MS2bs* [49] and *MAPT-MS2bs* [50] minigenes were gifts from Dr. Adrian Krainer (Cold Spring Harbor Laboratory, USA) and Dr. Gene Yeo (University of California, USA), respectively. HeLa (*SMN2-MS2bs*) or HEK-293T (*MAPT-MS2bs*) cells were plated in 12-well plates and, at 70% confluence, co-transfected with 200-600 ng of FLAG- or MCP-tagged constructs (adjusted depending on relative expression levels) and 100 ng of the minigene vector. Cells were collected 48 h post-transfection, washed with PBS, and split for western blotting (relative protein expression) and RNA extraction followed by RT-PCR (exon inclusion). Total RNA was treated with DNase I (ThermoFisher Scientific, 30 min, 37°C) before reverse transcription to avoid plasmid DNA contamination, and PCR products were resolved on 12% acrylamide gels as described above.

### Orthogonal organic phase separation

Sarcoma cells were cultured in 15 cm dishes until reaching ∼80% confluence for each condition (non-crosslinked, UV-crosslinked and DSG+UV-crosslinked), then washed twice with cold PBS. In each experiment, cells corresponding to the DSG+UV condition were then incubated in PBS with 2 mM DSG for 30 min under gentle agitation at room temperature, then washed twice with PBS and UV-crosslinked with 400 mJ/cm2 at 254 nm (Biolink™ BJX, Vilber). In parallel, the UV-only condition was irradiated in the same conditions, while the non-crosslinked control was kept at 4°C. Cells were then washed once with PBS, lysed in 1 mL of TriPure™ (Sigma) and stored at −80°C.

The OOPS experiment was conducted as previously described [51] for each sample with minor modifications. For “RNase-positive” samples, RNA was then degraded by fifteen 30-s sonication/rest cycles using the high setting, incubation at 95°C for 5 min, and simultaneous RNase A (final concentration 10 µg/mL, ThermoFisher Scientific) and RNase T1 (final concentration 10 U/mL, ThermoFisher Scientific) treatments at 37°C for 4 h. Finally, RNA-bound proteins were recovered by precipitation of the final organic phase using methanol, resuspended in Tris-HCl buffer (100 mM pH = 8.5) complemented with Laemmli buffer and subjected to western blotting with equivalent fractions of total lysates.

### Biotinylated isoxazole precipitation

For ectopic expression experiments, HeLa cells were seeded in 10-cm dishes, transfected at 70% confluence with 5 µg of pDEST-3xFLAG plasmid encoding the protein of interest, and collected 48 h later. For sarcoma cell experiments, cells were seeded in 15-cm dishes for each condition (with or without RNase treatment) and collected at 80-90% confluence. Cells were lysed for 30 min at 4°C in 2 mL lysis buffer (20 mM Tris-HCl pH 7.5, 150 mM NaCl, 5 mM MgCl2, 0.5% NP-40, 10% glycerol, 0.5 mM PMSF, 20 mM β-mercaptoethanol) and centrifuged at 16,000 g for 10 min; a 5% input was collected from the supernatant. For sarcoma cell lysates, one half was then incubated with 10 µg/mL RNase A for 1 h at 37°C and the other kept at 4°C to prevent RNA degradation. Lysates were split into four conditions and incubated with 0, 10, 30 or 100 µM b-isox for 1 h at 4°C on a rotating wheel, then centrifuged at 16,000 g for 10 min; 50 µL of each supernatant was collected as the non-precipitated fraction, and pellets were washed once with 400 µL lysis buffer before a further 5 min centrifugation at 16,000 g. Pellets and supernatants were resuspended in Laemmli buffer and analyzed by western blotting with equivalent fractions of total lysate.

### *Gaussia* luciferase protein complementation assay

HEK-293T cells were plated in 24-well plates and co-transfected with 200-600 ng (adjusted depending on relative expression levels) at ∼70% confluence with each complementary pDEST-GLuc plasmid per well. Empty GLucN1/N2 vectors were co-transfected with the corresponding tagged protein as negative controls. Cells were collected 48 h post-transfection and lysed in 200 µL *Renilla* Luciferase Assay Lysis Buffer (*Renilla* Luciferase Assay System, Promega) with vigorous agitation for 25 min, and lysates were dispensed into 96-well white flat-bottom plates in technical triplicate. Luminescence was measured after injection of the luciferase substrate (*Renilla* Luciferase Assay Reagent) on a Tristar^2^S LB942 luminometer (Berthold Technologies; 1 s delay, 10 s integration). Normalized luminescence ratios (NLR) were calculated as the luminescence of the tested interaction divided by the sum of the two corresponding negative controls.

### Subcellular fractionation

HeLa cells were cultured in 10-cm dishes, transfected at 70% confluence with 5 µg of vectors encoding FLAG-tagged wild-type or mutant fusions, and collected 48 h later by trypsinization and centrifugation (350 g, 5 min, 4°C). Pellets were lysed on ice for 5 min in PBS with 0.08% NP-40 and 1X cOmplete protease inhibitor mix (Roche). An aliquot of the whole lysate was kept as the total input fraction. The remaining lysate was centrifuged at 10,000 rpm for 30 s at 4°C. The supernatant was collected as the cytoplasmic fraction, and the pellet was washed twice on ice with lysis buffer to yield the nuclear fraction. All three fractions were boiled in Laemmli buffer and analyzed by western blotting.

### *In vitro* transcription, RNA biotinylation and RNA pulldown

Sequences cloned into pcDNA3.1 downstream of a T7 promoter were transcribed in vitro from 1 µg of linearized template using the HiScribe® T7 High Yield RNA Synthesis kit (New England Biolabs, standard NTP mix). Fragment size was verified on 1% agarose gels. Five µL of RNA was biotinylated with the Pierce™ RNA 3’ End Biotinylation kit (ThermoFisher Scientific) and quantified by NanoDrop. For RNA pulldowns, Dynabeads™ MyOne™ Streptavidin C1 (Invitrogen) were washed twice in lysis buffer (LB; 50 mM Tris-HCl pH 7.5, 0.5 mM EDTA pH 8, 0.5% NP-40, 10% glycerol, 120 mM NaCl), blocked with 15 µg/mL BSA and 10 µg/mL yeast tRNA (ThermoFisher Scientific), and incubated with 1 µg biotinylated RNA for 4 h. Sarcoma cells were grown to 80% confluence in 15-cm dishes (one per condition), washed twice with cold PBS, and lysed for 30 min at 4°C in 1 mL LB with cOmplete™ Protease Inhibitor Cocktail (Roche) and Halt™ Phosphatase Inhibitor Single-Use Cocktail (ThermoFisher Scientific). Lysates were sonicated (30 s, 30% amplitude), centrifuged at 5,000 g for 10 min (5% of lysate kept as input), and pre-cleared for 2 h with 10 µL beads on a rotating wheel at 4°C. Cleared lysates were split evenly and incubated overnight at 4°C with constant rotation with streptavidin beads alone, non-biotinylated RNA (negative controls), or biotinylated RNA-coupled beads. Beads were then washed twice with LB, four times with high-salt LB (500 mM NaCl), and once more with LB, before resuspension in 100 µL PBS. Samples were boiled in Laemmli buffer and analyzed by western blotting or tandem MS.

### Exonuclease-assisted mapping of protein-RNA interactions (ePRINT)

#### Experimental workflow

shA673-1C cells were treated with dox for 5 days in 15-cm dishes along with untreated controls to ∼80% confluence in biological duplicate. Cells were washed twice with PBS and UV-crosslinked (400 mJ/cm2) before harvesting. Non-crosslinked dishes were harvested in parallel as controls of crosslinking efficiency. A 5% input aliquot was analyzed by western blotting to confirm KD efficiency. Remaining cells were scraped in 1 mL PBS, pelleted (300 g, 5 min), and stored at −80°C. The ePRINT experiment was performed as previously described [39]. Final libraries were sequenced on a NovaSeq™ 6000 v1.5 platform (Illumina; 30M paired-end reads/sample) by the GIGA-Genomics platform (University of Liège).

#### Peak calling and analysis

ePRINT data processing was adapted from a previously described protocol [52]. Non-exonuclease-treated crosslinked RNA samples were used as input controls. Unique molecular identifiers (UMIs) were extracted with UMI-tools [53] with the barcode pattern set to NNNNNNNNNN. Illumina adapters were trimmed using Cutadapt [54], and reads were sorted using fastq-sort (fastq-tools package). Sorted reads were first aligned to RepBase sequences to filter out repetitive elements, and subsequently aligned to the GRCh38 genome assembly using STAR. Aligned reads were sorted and indexed using SAMtools [55], and deduplicated using UMI-tools. The quality of all FASTQ files was assessed using FastQC. Peak calling was performed using CLIPper with the GRCh38_v40 genome and an FDR threshold of 0.01. For each sample, peaks were sorted, filtered, and overlapping coordinates were merged using BEDTools [56]. Read coverage over peak regions was normalized to sequencing depth for all samples. ePRINT footprints were normalized to input read coverage. Reproducible peaks showing at least 1.5-fold enrichment over input (binomial test, p-val < 0.05) were retained as the final peak set for each condition. Data processing of peaks was performed in R using the *GenomicRanges* package. Peak sets were annotated using the annotatePeak function of the *ChIPseeker* package according to UCSC hg38 transcript annotations with annotation priorities in the following order: `“5UTR”, “3UTR”, “Exon”, “Intron”, “Promoter”, “Downstream”, “Intergenic“`. Overlaps with CLIP-seq data were performed with the `findOverlaps` function (*GenomicRanges*) between POSTARS3 CLIP regions and ePRINT peak ranges extended by 50 nt in both directions. Random peak coordinates of similar length were generated using BEDTools within human transcriptome intervals. Peak sequences were extracted from the UCSC hg38 genome using the *BSgenome* package. Sequences were shuffled using the *Biostrings* package. Peaks were defined as condition-specific when their enrichment was significant in one condition and not the other, or when the enrichment was at least two-fold higher in one condition than the other. They were otherwise defined as shared between both conditions and peak coordinates were merged using BEDTools. Peak relevance was assessed relative to EWSR1::FLI1-dependent splicing events identified in shA673-1C cells. Comparative motif enrichment analysis was performed using AME (MEME suite) with default parameters.

### Proteomic analysis of EwS cells

#### Sample preparation

shA673-1C, SK-N-MC/TR/shEF1, and MHH-ES-1/TR/shEF1 cells were seeded in 15-cm dishes at 50% confluence and treated with dox for 5 days along with non-treated controls. Cells were washed three times with PBS and harvested by scraping. A 5% whole-cell-extract aliquot was collected for western blotting to confirm KD efficiency. Cells were then pelleted (800 g, 5 min) and stored at −80°C.

Pellets were washed twice in ice-cold PBS and lysed in 100 µL lysis buffer containing 1% sodium deoxycholate (SDC), 10 mM TCEP, 40 mM chloroacetamide and 100 mM Tris (pH 8.5). Lysates were incubated on ice for 20 min, boiled at 95°C for 10 min, and sonicated for 20 min using a Bioruptor as previously described [57]. Lysates were digested with Trypsin and LysC (1:100 enzyme-to-protein ratio) overnight at 37°C with shaking at 1200 rpm. Digestion was stopped with five volumes of isopropanol containing 1% trifluoroacetic acid (TFA). Two layers of styrene-divinylbenzene reversed-phase sulfonate (SDB-RPS) stagetips were equilibrated with 100 µL of 30% methanol/1% TFA and 100 µL of 0.2% TFA; 20 µg of lysate was loaded, washed with 200 µL isopropanol/1% TFA and 0.2% TFA, and eluted with 60 µL of 80% acetonitrile (ACN)/1.25% ammonium hydroxide. Peptides were dried at 45°C for 90 min using a speed vacuum concentrator, re-diluted in buffer A* (2% ACN, 0.1% TFA), quantified by NanoDrop, and stored at −20°C.

#### Data-dependent acquisition (DDA) measurement

Peptides (400 ng) were loaded onto a nanoElute system coupled to a TIMS TOF HT mass spectrometer (Bruker Daltonics, Bremen, Germany) via a CaptiveSpray nano-electrospray ion source. Separation was performed on an IonOpticks Aurora 25 cm column using a binary solvent system: buffer A (0.1% formic acid, 2% ACN) and buffer B (0.1% formic acid in 99.9% ACN). Peptides were separated using a 120-min gradient starting at 2% buffer B, increasing to 12% in 60 min, 20% in 30 min, 30% in 10 min, and finally 85% in 10 min and held for an additional 10 min with a flow rate of 0.3 µL/min. TIMS elution voltage was calibrated using Agilent ESI-Low Tuning Mix ions (m/z 622, 922 and 1222) with known reduced ion mobility coefficients (1/K0). All measurements were conducted in dda-PASEF mode with a recording window from m/z 100 to 1,700 and a dimension range of 1/K0 = 0.65-1.40. Each top N acquisition cycle included 10 PASEF MS/MS scans. Precursor ions were selected for MS/MS based on an intensity threshold of 1,000 a.u. and re-sequenced until reaching a threshold of 10,000 a.u. TIMS operated within a scan range of 100–1,700 m/z, ramp time 100 ms, duty cycle 100%, cycle time 100 ms, and spectra acquisition rate 9.43 Hz.

#### Data processing

Raw DDA MS data were processed using FragPipe. Spectra were searched against the Homo sapiens UniProt reference FASTA supplemented with sequences derived from EWSR1::FLI1-dependent AS events translated in all three reading frames. Default LFQ Match-between-runs (MBR) workflow was applied. Trypsin was set for enzymatic cleavage, allowing up to 2 missed cleavages. The precursor mass tolerance was set to ±20 ppm and the fragment mass tolerance to ±20 ppm. Peptide lengths were restricted to 7–50 amino acids and precursor mass range to 500–5,000 Da. Label-free quantification was carried out using IonQuant with MaxLFQ enabled. Quantification was based on unique razor peptides, with at least one ion required for quantification. Peptide and protein identifications were filtered to a 1% false discovery rate. Protein and peptide differential expression were assessed in R using the *MSstats* package with default settings [58]. Proteomic data from the DSRCT cell lines JN-DSRCT-1/TR/shWT1 and SK-DSRCT-2/TR/shWT1 were previously described in [12], and were reprocessed with *MSstats* for differential protein expression analysis.

### Proteomic analysis of RNA pulldown lysates

Proteins were separated on a 10% SDS-PAGE gel with a short 15 min run at 120 V, visualized by colloidal Coomassie Blue staining, and in-gel digested with trypsin. Peptides were extracted with 0.1% TFA in 65% ACN and dried in a SpeedVac. Peptides were dissolved in solvent A (0.1% TFA in 2% ACN), directly loaded onto reversed-phase pre-column (Acclaim PepMap 100, ThermoFisher Scientific) and eluted in backflush mode. Peptide separation was performed using a reversed-phase analytical column (Acclaim PepMap RSLC, 0.075 x 250 mm, ThermoFisher Scientific) with a linear gradient of 4%-27.5% solvent B (0.1% FA in 98% ACN) for 40 min, 27.5%-50% solvent B for 20 min, 50%-95% solvent B for 10 min and holding at 95% for the last 10 min at a constant flow rate of 300 nL/min on a Vanquish Neo system. Peptides were analyzed by an Orbitrap Fusion Lumos tribrid or Exploris 240 mass spectrometer (ThermoFisher Scientific), then subjected to NSI source followed by tandem mass spectrometry (MS/MS) in Fusion Lumos or Exploris 240 coupled online to the nano-LC. Intact peptides were detected in the Orbitrap at a resolution of 120,000. Peptides were selected for MS/MS using HCD setting at 30, ion fragments were detected in the Orbitrap at a resolution of 30,000 (Exploris240) or in the linear ion trap (LUMOS). A data-dependent procedure that alternated between one MS scan followed by MS/MS scans was applied for 3 s for ions above a threshold ion count of 2.0E4 in the MS survey scan with 40.0 s dynamic exclusion. The electrospray voltage applied was 2.1 kV. MS1 spectra were obtained with an AGC target of 4E5 ions and a maximum injection time of 50 ms, and MS2 spectra were acquired with an AGC target of 5E4 ions and a maximum injection time set to dynamic. For MS scans, the m/z scan range was 375 to 1800. The resulting MS/MS data was processed using Sequest HT search engine within Proteome Discoverer 2.5 SP1 against the *Homo sapiens* protein database obtained from Uniprot. Trypsin was specified as cleavage enzyme allowing up to 2 missed cleavages, 4 modifications per peptide and up to 5 charges. Mass error was set to 10 ppm for precursor ions and 0.1 Da for fragment ions (LUMOS) or 10ppm (Exploris240). Oxidation on Met (+15.995 Da), conversion of Gln (−17.027 Da) or Glu (−18.011 Da) to pyro-Glu at the peptide N-terminal were considered as variable modifications. FDR was assessed using Percolator and thresholds for protein, peptide and modification site were specified at 1%. For abundance comparison, abundance ratios were calculated by Label Free Quantification (LFQ) of the precursor intensities within Proteome Discoverer 2.5 SP1. Subsequent data processing was performed in R.

### Cell proliferation

Twenty-four hours post-transfection, cells were detached, counted, and seeded into a 96-well plate in technical duplicates or triplicates at a density calculated to achieve approximately 30% confluence the following day. Two hours prior to the first timepoint, cell nuclei were stained using SPY650-DNA (1:2,000 in medium, Spirochrome). Phase as well as near-infrared fluorescent images were then acquired every 4 hours for 44 hours using the Incucyte SX5 Live-Cell Analysis System (Sartorius). Cell proliferation was quantified over time by automated nucleus counting (A-673, SK-DSRCT-2, MLS-402) or by automated confluence calculation (KAS) using the Incucyte SX5 integrated analysis software (Sartorius). Nuclei counts or confluence values were normalized to the first timepoint (T0).

### Cell viability

Twenty-four hours post-transfection, cells were detached, counted, and seeded in technical triplicates into a 96-well plate (Greiner). Forty-eight hours after seeding, cell medium was removed and replaced with resazurin working solution (medium with 1 µg/mL resazurin, Stemcell Technologies). Unseeded wells containing only the resazurin working solution were incubated under the same conditions to serve as blanks. Following incubation for 3 hours at 37°C, the medium containing metabolized resazurin was transferred to a black 96-well plate (Greiner) for fluorescence detection using a FilterMax F5 plate reader (Molecular Devices) at excitation and emission wavelengths of 535 nm and 595 nm, respectively. Measurements were normalized to the control condition (siCTL) following blank subtraction.

### RNA-sequencing analysis

When applicable, raw FASTQ data were retrieved from the NCBI Sequence Read Archive using the SRA toolkit. To enable data integration and ensure reproducibility, RNA-seq analysis was systematically performed using the nf-core/rnaseq v3.18 pipeline with FASTQ files as input and the singularity profile from the GIGA institute [59]. Briefly, sequencing adapters and reads of poor quality were trimmed with TrimGalore! and Cutadapt, and aligned to the GRCh38 human genome assembly using STAR [60]. Transcript assembly and quantification of aligned reads were performed with Salmon [61] in alignment-based mode and StringTie [62] according to Ensembl release 114 annotations. All quality controls performed throughout the pipeline were analyzed with MultiQC [63].

Differential gene expression analysis was performed using *DESeq2* [64] using a model including condition as the primary factor. PCA was performed on variance-stabilized counts in R with the `prcomp` function. 3D PCA plots were obtained with the *plotly* package. Genes were annotated using the *biomaRt* package.

### Alternative splicing quantification

Genome-aligned reads were used as input for alternative splicing analysis using rMATS-turbo [65] with the `--variable-read-length --novelSS` flags. Subsequent data processing was performed in R with output JCEC tables. For individual datasets, detected AS events were filtered based on FDR below 0.05, IncLevelDifference absolute values (referred to as PSI) greater than 0.05, and events supported by a mean ≥ 10 junction reads across replicates. For data integration of splicing results, events were merged according to identical chromosomal coordinates and filtered on FDR in overlapping datasets (FDR < 0.05).

### ChIP-sequencing analysis

ChIP-seq datasets of fusion binding were retrieved from previous studies [6,9,11,66,67,48,68,69]. When applicable, raw FASTQ files were retrieved using the SRA toolkit. ChIP-seq analysis was performed with the nf-core/chipseq v2.1 pipeline with FASTQ files using the `--narrow-peak` parameter and the singularity profile from the GIGA institute. Briefly, after trimming, reads were aligned to the GRCh38 using BWA [70]. Read duplicates were marked with Picard and high-confidence narrow peaks were called with MACS [71]. Within each experiment, overlapping peaks were merged across replicates or integrated between related cell lines. Data from EwS and DSRCT ChIP-seq were directly retrieved from Supplementary Tables of the original studies, and peak coordinates were converted to hg38 genome coordinates using R *liftOver* package.

Final peak sets were imported into R and annotated using the `annotatePeak` function of the *ChIPseeker* package and transcript annotations from the *TxDb.Hsapiens.UCSC.hg38.knownGene* package. Motif analysis was performed by calling the `findMotifsGenome.pl` wrapper from HOMER with the parameters `-size 200 -p 8`. Nearest genes were assigned to each peak based on its closest annotated protein-coding gene’s transcription start site. Overlaps with AS events were computed using the `findOverlaps` function (*GenomicRanges*) between peak ranges and event ranges extended up to 50 kb in each direction.

### Sequence properties of splicing events

Junction coordinates from alternative splicing events were used to build `GRanges` objects using the *GenomicRanges* R package. Genomic regions corresponding to these ranges were extracted using the *Biostrings* and *BSgenome.Hsapiens.UCSC.hg38* packages as `DNAStringSet` objects and exported to FASTA files. These files were used as input for splicing signal score computation. 5’ and 3’ splice site scores were determined with the MaxEntScan algorithm [72]. SVM-BPfinder was used to compute BP and PPT scores [73]. GC content was calculated using the `GC` function of the *Biostrings* package. For rG4 enrichment analysis, sequences spanning the region from the upstream to the downstream exon were scanned for the consensus motif using the regular expression `G{3,5}.{1,7}G{3,5}.{1,7}G{3,5}.{1,7}G{3,5}` for all detected events. Additionally, experimentally validated rG4 coordinates were retrieved from the QUADRatlas database [74] and intersected with splicing event coordinates to assess the average number of rG4s in each AS event set. Enrichment significance was assessed using an empirical p-value calculated from 10,000 random permutations of background region sets of similar size. To establish AS exon positions within the gene body, genomic coordinates of each exon were matched to Ensembl release 114 annotations to determine its exon rank within the longest transcript. If the SE could not be associated to previously annotated exons, the exon rank number was assigned as the upstream exon rank plus one. Relative position within the gene body was determined as the ratio between the SE rank and the total number of exons in that gene.

### Functional enrichment of gene sets

Enrichment analyses of gene sets from the Gene Ontology, KEGG and REACTOME databases were performed using the R package *clusterProfiler* and *ReactomePA.* The COSMIC Cancer Gene Census list was retrieved from the official COSMIC website [75]. Entrez gene identifiers were used for comparison with sets of differentially spliced genes from the four sarcoma landscapes.

### Prediction of transcript fate

Splicing graphs were reconstructed from exon-exon junction coordinates associated with AS events. For each event, two transcript sets (inclusion vs. exclusion) were defined according to GENCODE v48 annotations. To infer the predicted impact of differential exon inclusion on transcript fate, transcript biotypes were compared between the two sets. Events were classified as follows as “poison exons” if none of the exon-containing transcripts were annotated as protein-coding, at least one was annotated as nonsense-mediated decay, and at least one exon-skipping transcript was protein-coding. Events were classified as “essential exons” if at least one transcript including the exon was protein-coding, while transcripts lacking the exon were not protein-coding and at least one was annotated as nonsense-mediated decay. Alternatively, experimentally defined NMD-sensitive events were retrieved from the dataset of [33], which profiled splicing changes after UPF1/XRN1 co-depletion. Overlaps between landscape events and NMD-regulated events (p < 0.01) were computed using the `findOverlaps` function (*GenomicRanges*).

### RNA-binding protein motif and pentamer enrichment

A compendium of 836 RBP motifs was retrieved from the CISBP-RNA [76] and oRNAment [77] databases and uniformly converted into position frequency matrices. Production of RBP splicing maps was adapted from [78] and [79]. Sequences were systematically scanned for motif occurrences using a sliding-window algorithm (window size = 50, step = 1) across 400 nt regions spanning exon-intron junctions (300 nt intronic and 100 nt exonic sequences) with the `matchPWM` function from the *Biostrings* R package (minimal match score = 80%). For pentamer enrichment analysis, all 1,024 possible pentamers were scanned along the same sliding windows using the `matchPattern` function (*Biostrings*).

Motif enrichment was assessed in each sliding window using an FDR-corrected Wilcoxon rank-sum test comparing significant events to background events. To complement these approaches, *de novo* motif discovery was performed using DREME (MEME suite) [80].

Enriched pentamers and DREME-identified motifs (adjusted p-value < 0.05) were assigned to specific RBPs by matching similarity with position frequency matrices of the curated collection of 836 experimentally validated motifs. Motif similarity was evaluated using the *universalmotif* R package, and matches were retained when the Euclidian distance-based similarity score exceeded 0.8. For each RBP, enrichment significance was summarized using the lowest FDR in every junction among its associated motifs. To evaluate exon inclusion dependence on positional effects, motif enrichment was compared between the upstream and downstream exon-intron junctions using an FDR threshold of 0.01.

### RNA-binding protein binding site enrichment

CLIP-seq data were retrieved from the POSTAR3 database [81]. For each RBP, overlaps between AS event ranges and CLIP peak ranges were computed with the `findOverlaps` function (*GenomicRanges*) with default parameters. Enrichment significance was assessed using an empirical p-value calculated from 10,000 random permutations of background regions.

### ENCODE RNA-seq analysis

GRCh38 genome-aligned BAM files of RNA-seq analysis following RBP shRNA knockdown or CRISPR-Cas9 knockout experiments in K562 and/or HepG2 cells were retrieved from the ENCODE data portal. These files were used as input for rMATS alternative splicing analysis with the same parameters as described above.

### Motif co-occurrence

Motif co-occurrence analysis was performed using motif matches identified in the exon-intron junction regions (400 nt sequences) from the sliding-window enrichment analysis described above. For each pairwise motif comparison, contingency tables were constructed within each sequence set based on presence or absence of the two motifs. Log-transformed odds ratios (LOR) were computed to quantify co-occurrence, defined as the binary logarithm of the ratio between concordant observations (both motifs present or both absent) and discordant observations (only one motif present). To prevent inflated co-occurrence scores caused by motif redundancy, similarity between position frequency matrices was evaluated using the *universalmotif* R package. Highly similar motifs (distance score > 0.7 or p < 0.05) were excluded. Motif pairs with significant LOR values (above the 95th percentile of the distribution) were retained for downstream analysis. Only motifs previously identified as significantly enriched in the sliding window analysis were ultimately retained. Finally, motif co-occurrence was considered position-specific (5’ or 3’) when the LOR value was significant in one region but not the other, or when the LOR score in one region exceeded that of the opposite junction by at least twofold. Co-occurrence networks were analyzed and visualized with Cytoscape.

### Protein-protein interaction predictions

EWSR1::FLI1 and FUS::DDIT3 interaction networks were retrieved from their original publications [20,21]. Protein-protein interactions involving the 51 fusion-associated RBPs were retrieved from the HINT database of high-quality binary and co-complex interactions [37]. Additionally, this dataset was complemented with published interactome data for FUS [82], ARGLU1 [83], RBM25 [84], RBM45 [85], SRSF1 [86], RBM42 [87] and RBM22 [88]. Gene symbols were processed with R to filter duplicates and ensure a consistent gene nomenclature. Network visualization was performed using Cytoscape.

### Homology-based clustering of RBPs

Multiple sequence alignment was performed on all RBP sequences retrieved from the UniProt database using the CLUSTALW algorithm [89] with the BLOSUM62 matrix (gap opening penalty = 10, gap extension penalty = 1). Phylogenetic trees were constructed using the *ape* R package.

### EwS cohort transcriptomic analysis

EwS patient clinical metadata and transcriptomic data were retrieved from the BOCA-FR cohort (data portal release 28) of the International Cancer Genome Consortium (ICGC) [90]. Raw sequencing files were downloaded from the ICGC controlled-access portal using the pyEGA3 client. FASTQ files were used as input for the Nextflow nf-core/rnaseq pipeline. Exon inclusion levels (PSI) were quantified using *outrigger* on genome-aligned BAM files with the corresponding annotation files and default parameters. PSI matrices were filtered to include events detected in at least 80% of patients. Missing values were imputed with the *VIM* R package using the k-nearest neighbor classification.

Survival analyses were performed in R using the *survival* and *survminer* packages. The multivariate Cox proportional hazards regression analysis was performed using the coxph function with default parameters. One patient with unknown tumor stage at time of diagnosis was excluded from the multivariate model. For each event, PSI values were stratified into three groups by k-means clustering. Data integration with landscape events was obtained by reconstructing splicing graphs and matching exon-exon junction coordinates to events filtered by *outrigger*.

### Data visualization and statistics

Unless otherwise specified, all data processing was performed in R (RStudio environment). Overlap significance was computed using the *GeneOverlap* R package. Statistical testing of experimental results was performed using GraphPad Prism or in R. Phylogenetic and clustering trees were visualized from exported dendrograms (R package *dendextend*) with iTOL [91]. Depth-normalized bigwig files of aligned reads were used for visualization using IGV [92]. Gel images were processed with ImageJ/Fiji. Graphs were generated using GraphPad Prism. Schematic figures were constructed using BioRender and Adobe Illustrator. All software and packages are listed in **Supplementary Table S3**.

### Data availability

Custom scripts developed for this study are available at https://github.com/Dequiedt-lab/Fusion_Splicing_Scripts/. All datasets used in this work are listed in **Supplementary Table S3**. Newly generated sequencing data are *to be deposited*.

---

**Figure S1.**
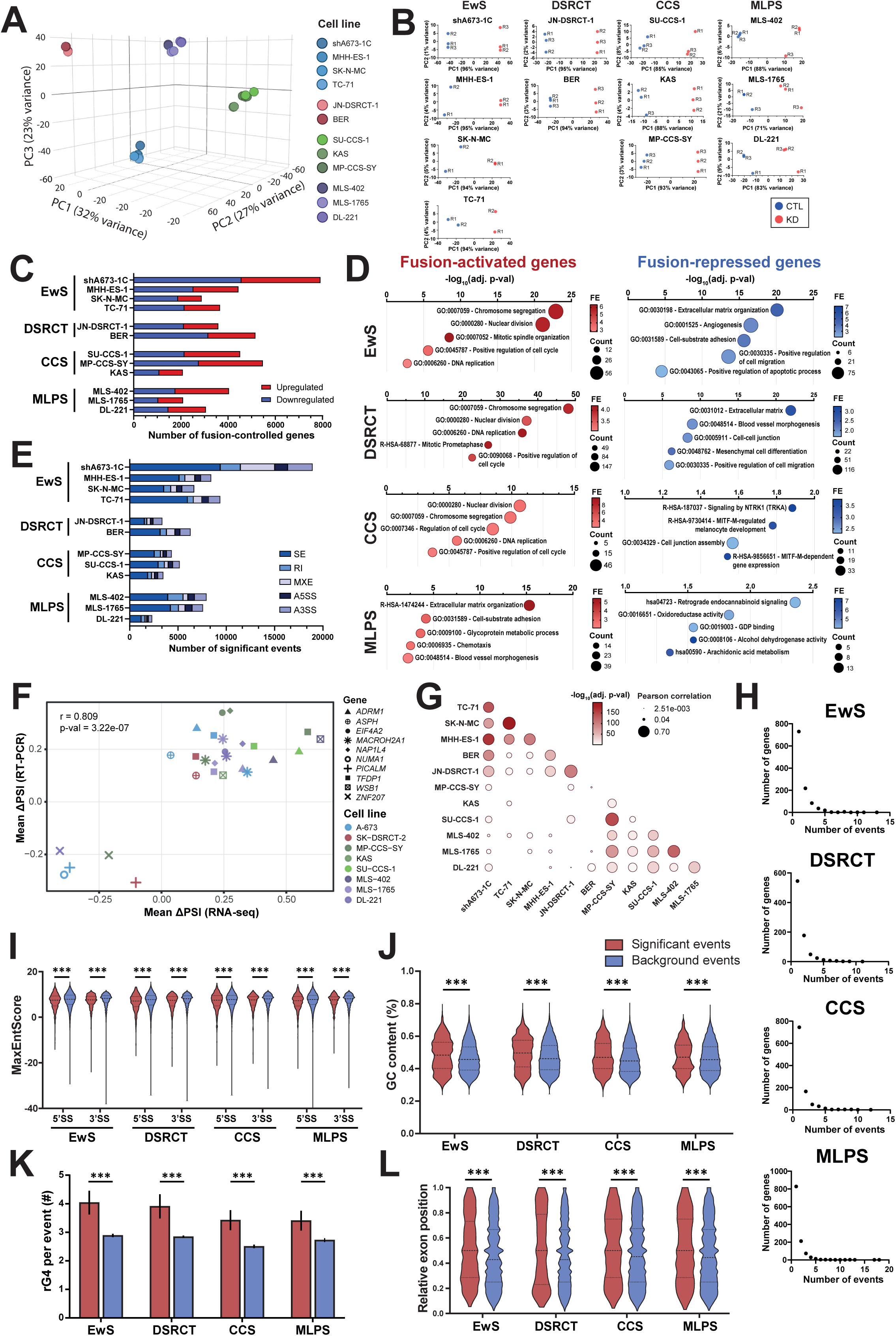
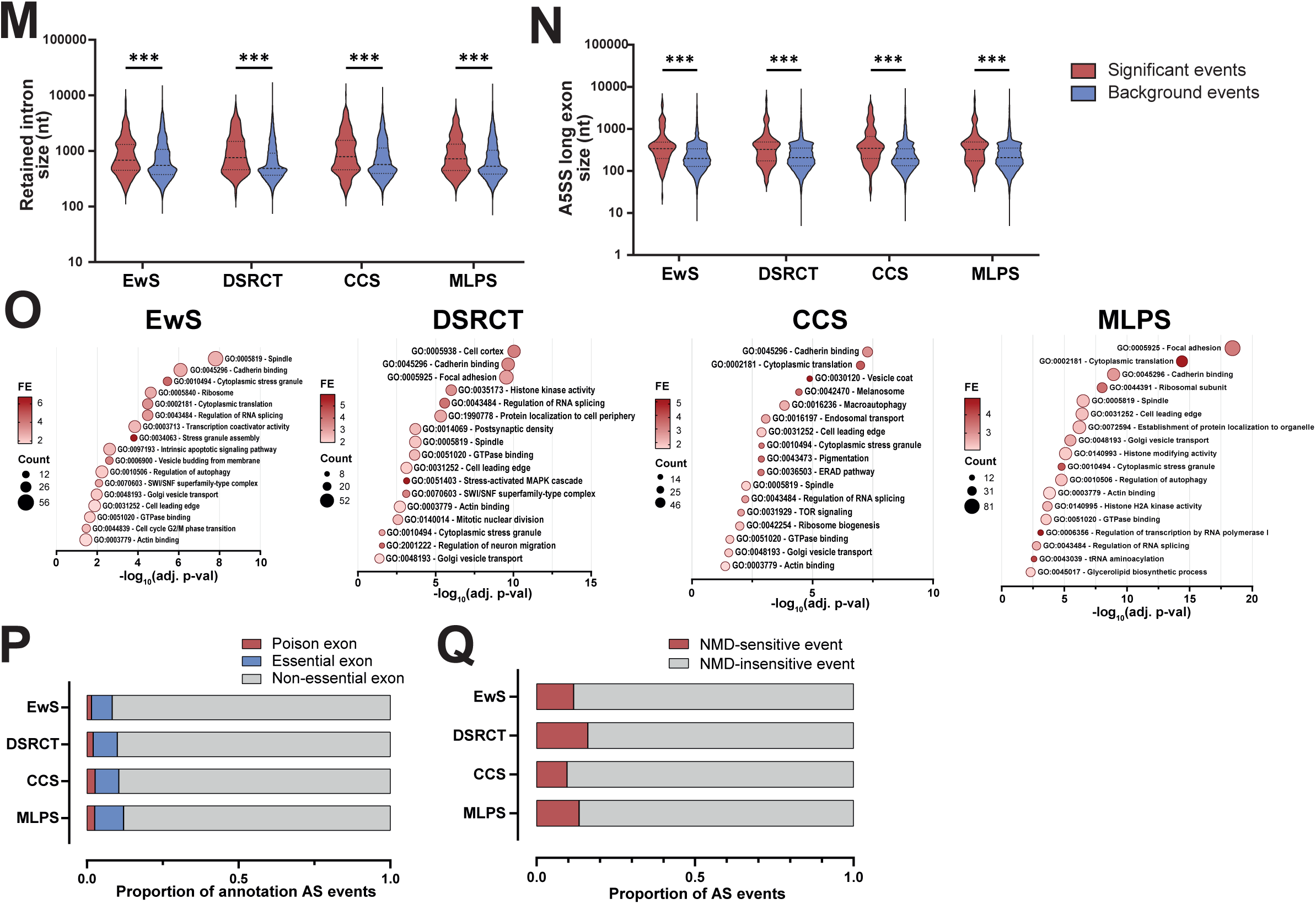
(**A**) Principal component analysis of basal gene expression profiles across the 12 sarcoma cell lines. (**B**) Principal component analysis of gene expression profiles in control vs knockdown (KD) conditions for each sarcoma cell line. (**C**) Differential gene expression in each FET-rearranged sarcoma cell line following fusion KD (|log_2_FC| > 1, FDR < 0.05). (**D**) Gene ontology and pathway terms enriched for genes activated and repressed by FET fusions. P-values were adjusted using the Benjamini-Hochberg method. FE: fold-enrichment. (**E**) Significant differential splicing events following fusion KD for each sarcoma cell line (FDR < 0.05, |ΔPSI| > 0.05, and mean junction counts > 10). (**F**) RT-PCR validation of fusion-dependent AS events. Validation was conducted on at least three independent biological replicates prepared under similar conditions to those used for RNA-seq samples. r: Pearson correlation coefficient. (**G**) Correlation matrix plot of significant splicing events between sarcoma cell lines. Pairwise Pearson correlations were computed between each pair of datasets with commonly detected skipped exon events. (**H**) Number of events per differentially spliced genes in the four splicing landscapes. (**I**) Splice site (SS) strength of the 5’SS and 3’SS of skipped exons for significant and background (FDR > 0.05 in all related cell lines) events across the four sarcoma landscapes. (**J**) GC content percentage around exon-intron junctions across the four sarcoma landscapes. (**K**) Number of experimentally validated RNA G-quadruplex (rG4) overlapping significant and background splicing events across the four sarcoma landscapes. (**L**) Distribution of exon positions along the gene body of fusion-regulated (significant) and background exons for the four sarcoma landscapes. (**M**) Sizes of retained introns in nucleotides (nt) for fusion-regulated (significant) and background events across the four sarcoma landscapes. (**N**) Sizes of alternative 5’ splice sites (A5SS) in nucleotides (nt) for fusion-regulated (significant) and background events across the four sarcoma landscapes. (**O**) Gene ontology enrichment analysis of differentially spliced genes from the EwS, DSRCT, CCS and MLPS landscapes. P-values were adjusted using the Benjamini-Hochberg method. FE: fold-enrichment. (**P**) Fraction of AS events categorized as potentially related to nonsense-mediated decay (NMD), i.e. essential or poison exons, or non-essential based on transcript annotations. (**Q**) Fraction of AS events that overlapped with events experimentally demonstrated to be associated with NMD.

**Figure S2.**
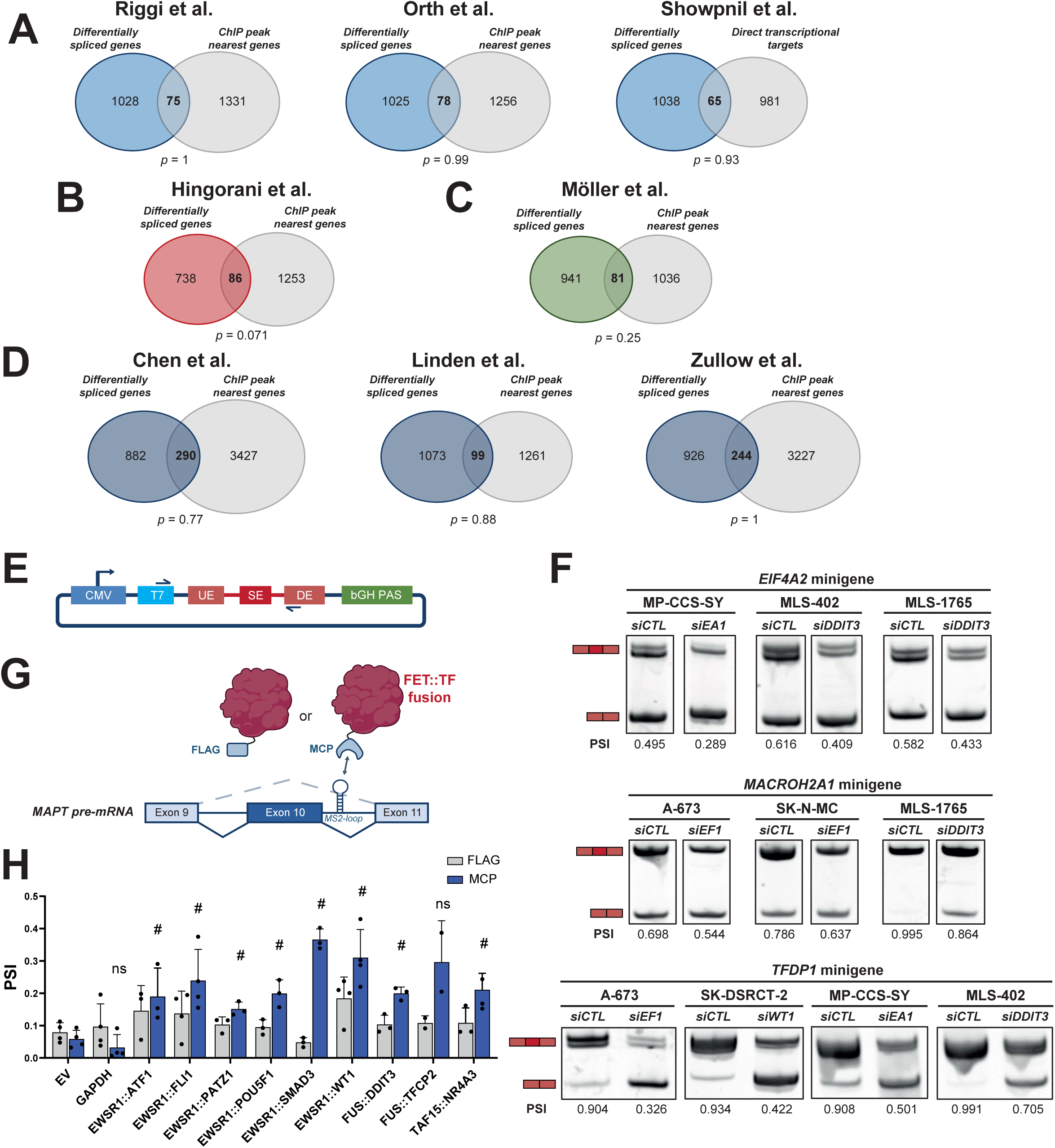
Overlap between differentially spliced genes and (**A**) genes associated with EWSR1::FLI1 chromatin binding and EWSR1::FLI1 direct transcriptional targets (as defined in [11]), (**B**) genes associated with EWSR1::WT1 chromatin binding, (**C**) genes associated with EWSR1::ATF1 chromatin binding, and (**D**) genes associated with FUS::DDIT3 chromatin binding. *p*: overlap p-value (Fisher’s exact test). (**E**) Schematic representation of the pcDNA3.1 vector with the inserted minigene sequence that spans the region from the upstream to the downstream exon of the splicing event. (**F**) Representative RT-PCR results of cassette exon inclusion levels upon fusion KD are depicted for the *EIF4A2*, *TFDP1* and *MACROH2A1* minigenes in multiple sarcoma cell lines. (**G**) Schematic representation of the MS2-tethering splicing assay of the *MAPT* reporter minigene. (**H**) Exon inclusion of the SMN2 minigene reporter upon transfection of FLAG or MCP-tagged empty vector (EV) and GAPDH controls, and a representative panel of 8 different FET fusions. ns: not significant; #: p-value < 0.1 (FDR-adjusted Wilcoxon–Mann–Whitney test).

**Figure S3.**
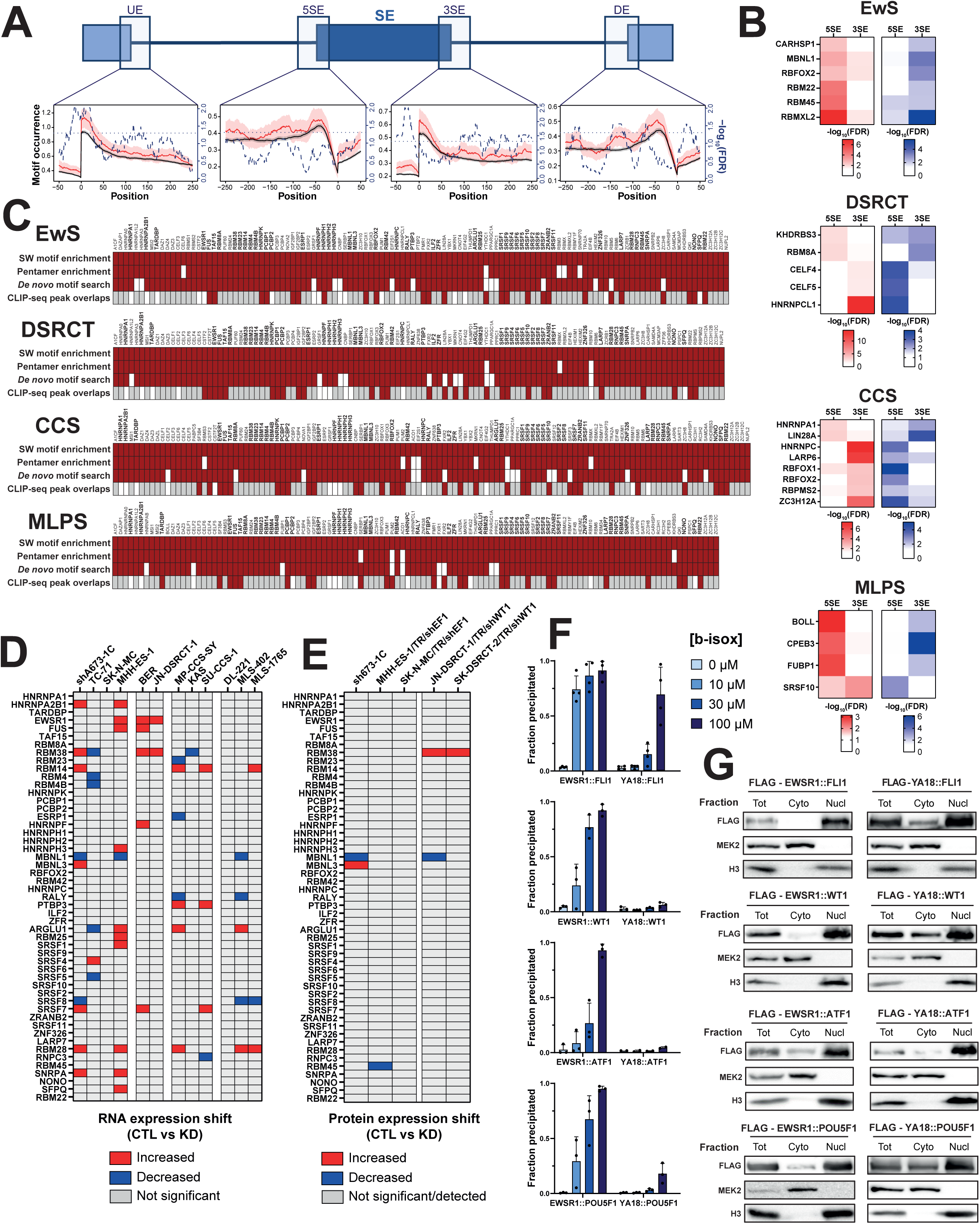
(**A**) Representative illustration of RNA splicing maps for RNA-binding protein motifs using a sliding-window algorithm. For each window corresponding to the four splicing junctions, motif occurrence is computed for significant AS events (represented as the plain red line) and tested against a background set of sequences (plain black line). Dashed blue lines represent the FDR-corrected p-values (log-transformed, Wilcoxon–Mann–Whitney test) scaled on the right y-axis. The dotted line marks the significance threshold. (**B**) Motif enrichment analysis of positively (ΔPSI > 0; red) and negatively regulated (ΔPSI < 0; blue) exons around the 5’ (5SE) and 3’ (3SE) splicing junction surrounding the cassette exon in the EwS, DSRCT, CCS and MLPS landscapes. FDR values inferior to 0.05 are not colored. (**C**) Summary of four enrichment analyses of RBP binding around fusion-regulated exons. RBPs with significantly enriched motifs in the sliding-window (SW) based analysis were cross-validated by random nucleotide pentamer enrichment analysis, *de novo* motif discovery using DREME, and the enrichment of CLIP-seq peaks around fusion-regulated events across the four splicing landscapes. Red rectangles mark significant enrichment. Gray rectangles indicate the absence of CLIP-seq data. The 51 splicing factors validated by at least two computational methods in every landscape are highlighted in bold. (**D**) Differential RNA expression of fusion-associated RBPs (|log_2_FC| > 1, adjusted p-value < 0.05) following fusion KD in the twelve sarcoma cell lines. (**E**) Differential protein expression of fusion-associated RBPs (|log_2_FC| > 1, adjusted p-value < 0.05) following fusion KD in EwS cells and two DSRCT cell lines (from [12]). (**F**) Comparative precipitation of wild-type and tyrosine mutant (YA18) fusions with increasing concentrations of biotinylated isoxazole (b-isox) for EWSR1::FLI1, EWSR1::POU5F1, EWSR1::WT1 and EWSR1::ATF1 following ectopic expression in HeLa cells. (**G**) Cytoplasmic and nuclear distribution of wild-type and YA18 fusions for EWSR1::FLI1, EWSR1::POU5F1, EWSR1::WT1 and EWSR1::ATF1 following ectopic expression in HeLa cells. MEK2 and H3 were used as cytoplasmic and nuclear protein markers, respectively.

**Figure S4.**
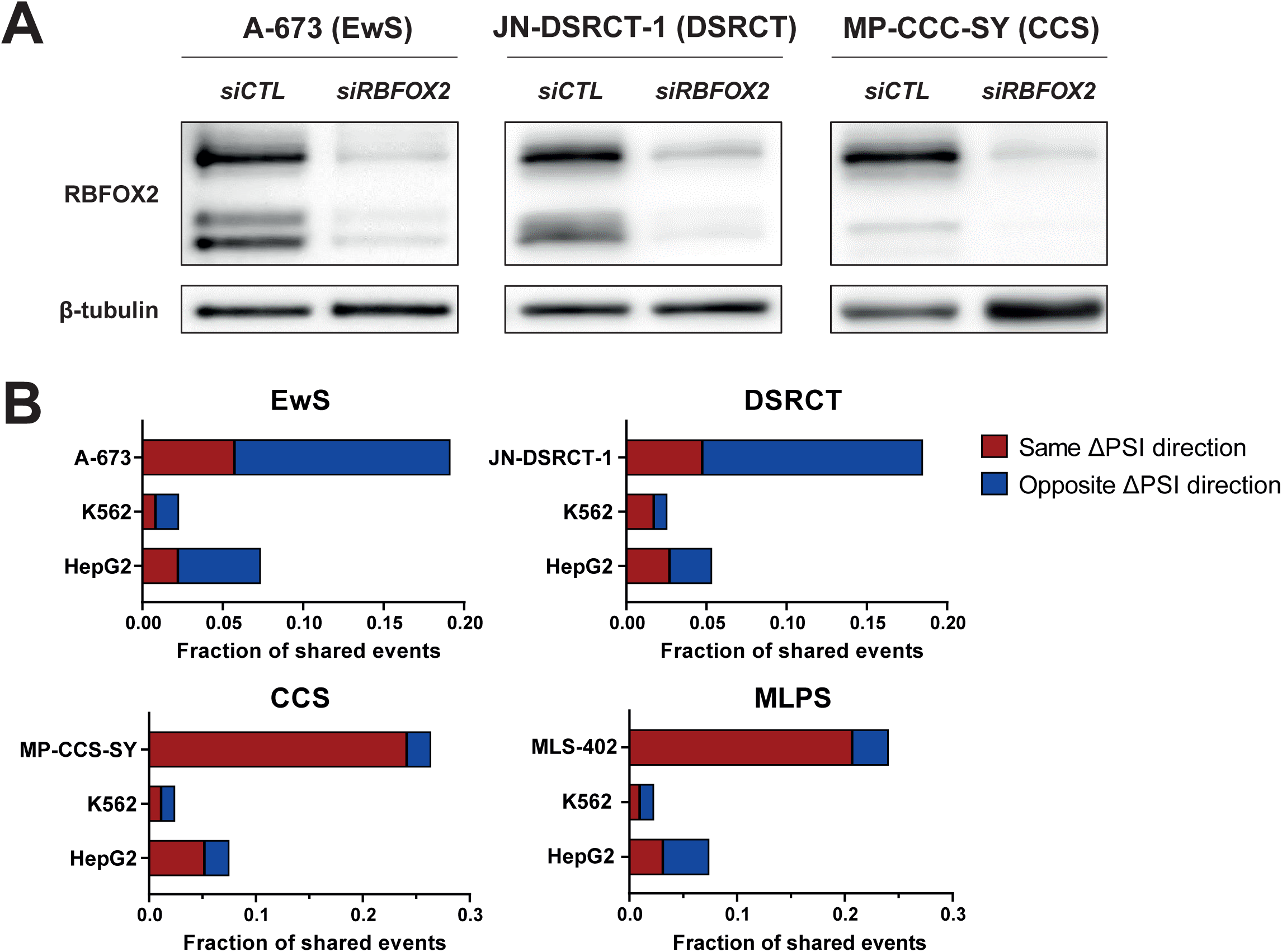
(**A**) Representative western blot of RBFOX2 levels in RNA-seq samples of sarcoma cells treated with control and RBFOX2-targeting siRNAs. β-tubulin was used as a loading control. (**B**) Comparison of RBFOX2 co-regulated events in K562, HepG2 (ENCODE datasets) and the appropriate sarcoma cell line for EwS, DSRCT, CCS and MLPS splicing landscapes.

**Figure S5.**
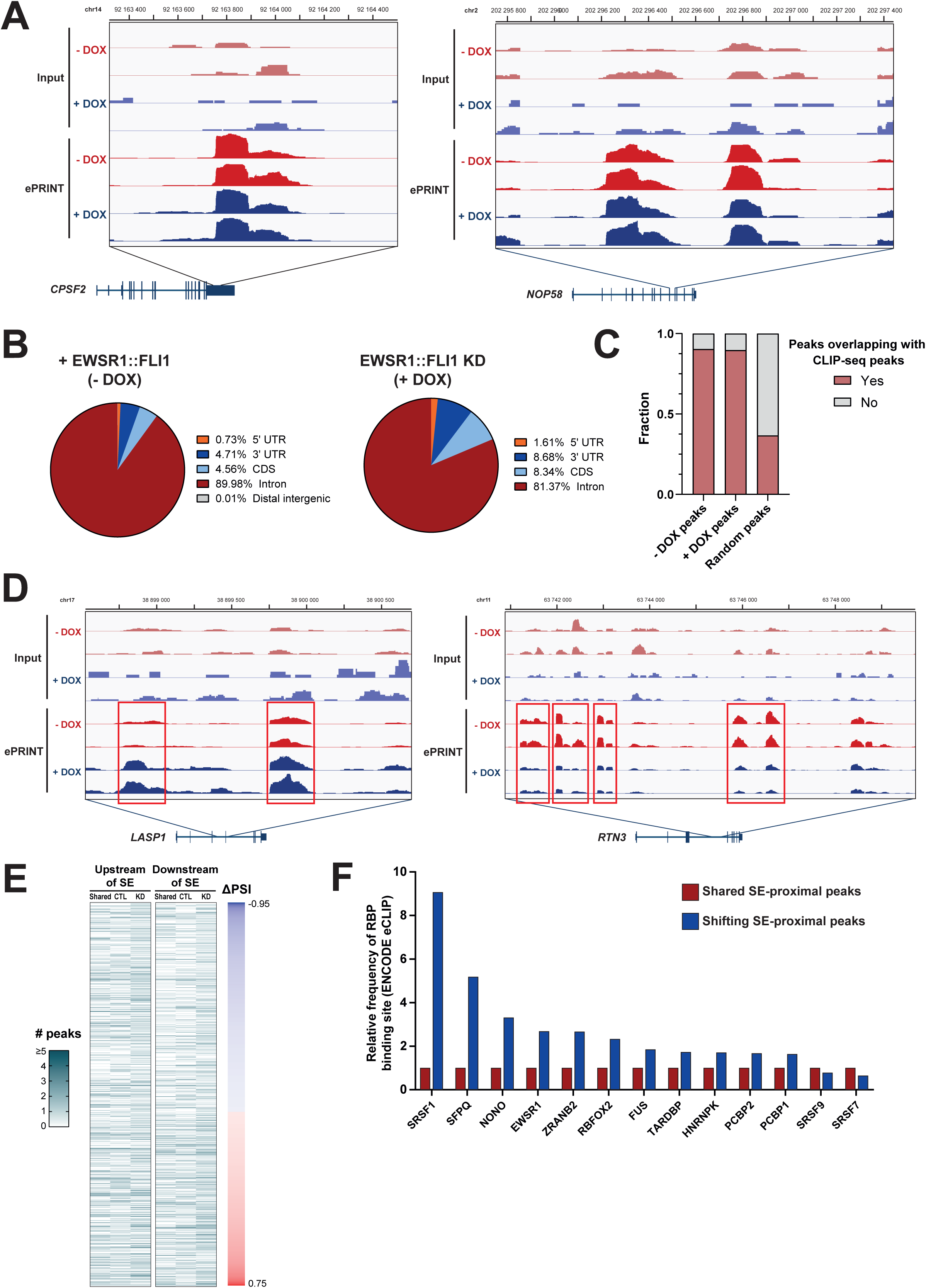
(**A**) Representative visualization of ePRINT peaks in the 3’UTR of *CPSF2* and an intronic segment of *NOP58*. (**B**) ePRINT peak location in the control (-DOX) and KD (+DOX) conditions. (**C**) Percentage of ePRINT peaks that overlapped with at least one previously reported CLIP-seq peak. (**D**) Representative examples of differential ePRINT peaks in control and fusion knockdown conditions. ePRINT tracks are shown for peaks dependent on EWSR1::FLI1 expression in intronic regions of the *RTN3* and *LASP1* genes. (**E**) Relationship between location and number of shared/shifting ePRINT peaks and exon inclusion levels. Number (#) of peaks was capped at 5. CTL: control-specific (-DOX) peaks. KD: knockdown-specific (+ DOX) peaks. (**F**) Relative prevalence of ENCODE eCLIP peaks overlapping with shared and shifting ePRINT peaks in proximity to EWSR1::FLI1-dependent cassette exons for available fusion partner RBPs.

**Figure S6.**
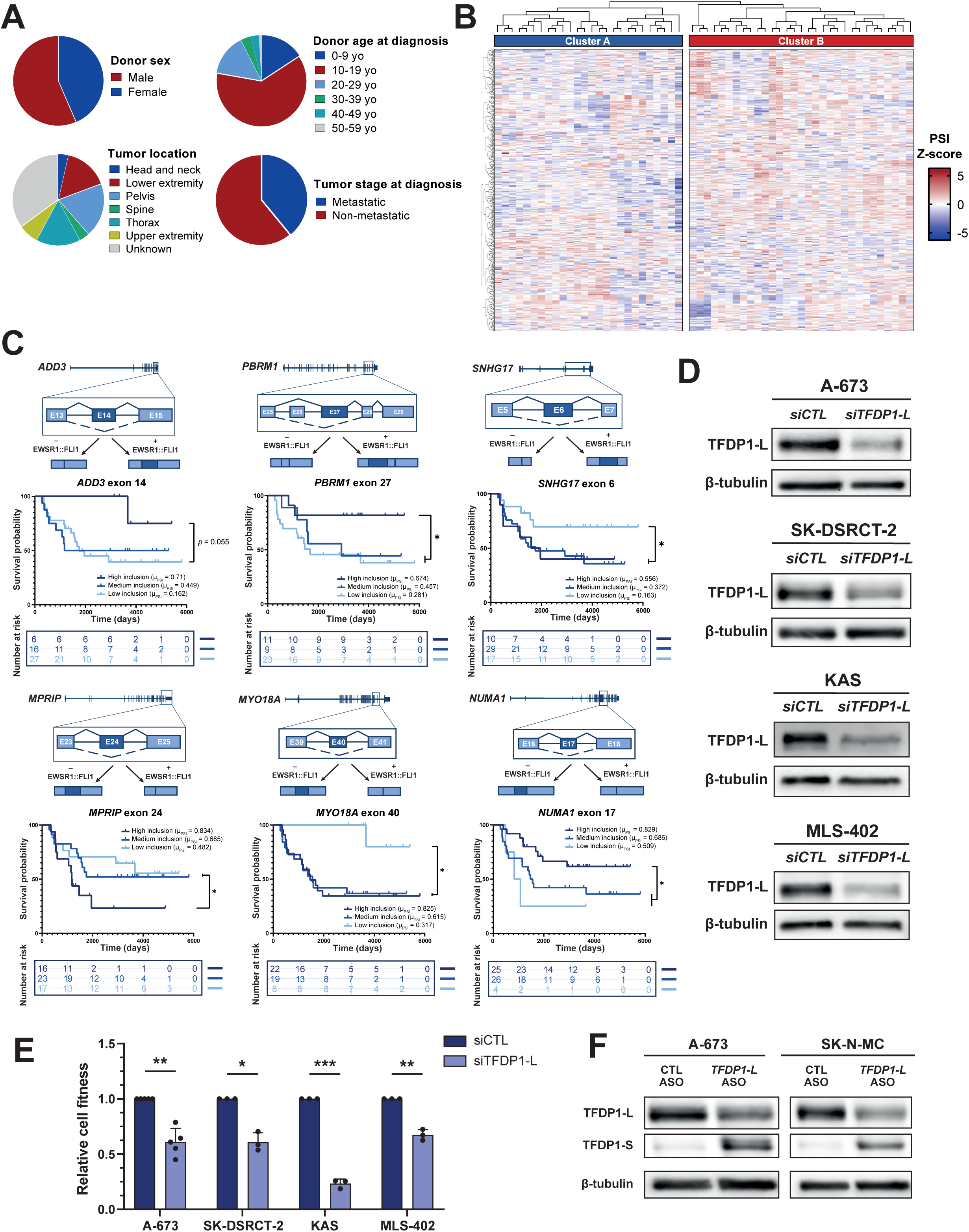
(**A**) Clinical features of the EwS patient cohort (n = 57). (**B**) Splicing event inclusion levels of the EwS patient cohort grouped into two distinct clusters (A and B) by unsupervised hierarchical clustering. For visualization, PSIs were z-score transformed and grouped via hierarchical clustering. Only the 500 events exhibiting the highest variance between patient samples are represented. (**C**) Kaplan-Meier estimation curves for a panel of representative survival-associated EWSR1::FLI1-dependent AS events in the *ADD3*, *PBRM1*, *SNHG17*, *MPRIP*, *MYO18A* and *NUMA1* genes. *: p-value < 0.05 (log-rank Mantel-Cox test). (**D**) Representative western blot of TFDP1-L protein levels 72 h following transfection of an isoform-specific siRNA (*siTFDP1-L*) in A-673, SK-DSRCT-2, KAS, and MLS-402. (**E**) Relative cell viability of A-673, SK-DSRCT-2, KAS and MLS-402 cells 72 h following transfection of *siTFDP1-L*. Results are displayed as means of 3 or more individual biological replicates normalized to the control condition (siCTL). *: p-value < 0.05; **: p-value < 0.01; ***: p-value < 0.001 (one-sample t-test) (**F**) Representative western blots of the long (TFDP1-L) and short (TFDP1-S) TFDP1 protein isoforms in EwS cells 72h following ASO treatment. 200 nM ASO and 100 nM ASO were transfected in A-673 and SK-N-MC cells, respectively. β-tubulin was used as a loading control.

**Table 1.** Multivariate Cox proportional hazards regression analysis of overall survival. Hazard ratios (HR) and 95% confidence intervals (CI) were estimated for patient cluster assignment, donor sex, tumor stage at diagnosis and age at diagnosis. Reference categories are indicated in parentheses.

| Variable | HR | 95% CI | z | p-value |
| --- | --- | --- | --- | --- |
| Patient cluster ("B" vs. "A") | 2.904 | 1.317–6.404 | 2.647 | 0.008 |
| Sex (male vs. female) | 1.042 | 0.491–2.213 | 0.108 | 0.914 |
| Tumour stage (non-metastatic vs. metastatic) | 0.620 | 0.283–1.359 | -1.194 | 0.232 |
| Age at diagnosis | 0.951 | 0.888–1.019 | -1.414 | 0.157 |

